# Diversity and ecological functions of viruses inhabiting the oil reservoirs

**DOI:** 10.1101/2023.09.19.558540

**Authors:** Liyun An, Xinwu Liu, Jianwei Wang, Jingbo Xu, Xiaoli Chen, Xiaonan Liu, Bingxin Hu, Yong Nie, Xiao-Lei Wu

## Abstract

Oil reservoirs, being one of the most significant subsurface repositories of energy and carbon, have long hosted diverse microorganisms affecting energy production and carbon emissions. Viruses play crucial roles in the ecology of microbiomes, however, their distribution and ecological significance in oil reservoirs remain undetermined. Here, we assembled an extensive catalogue encompassing viral and prokaryotic genomes sourced from oil reservoirs. The catalogue comprises 7,229 prokaryotic genomes and 6,218 viral genomes from 182 oil reservoir metagenomes, respectively. Based on sequence clustering we identified 3,886 Operational Taxonomic Units (vOTUs) approximately at species level, 94.80% of which were not found in any other environments. The results showed that viruses were widely distributed in oil reservoirs, and oil reservoirs contain a significant number of unique and unexplored viruses. We also constructed a catalogue of 322,060 viral gene clusters including 105 virus-encoded putative auxiliary metabolic genes (AMGs) that participate in host metabolism and adaptation to the environment. Furthermore, our investigation has yielded a total of 7,197 putative virus-host pairs based on CRISPR and tRNA profiles. Viruses within oil reservoirs profoundly infected bacterial Actinobacteriota, Desulfobacterota, as well as archaeal Halobacteriota and Methanobacteriota. Combined microcosm enrichment experiments and bioinformatics analysis, we validated the ecological roles of viruses in regulating the community structure of sulfate reduction microorganisms, primarily through the predation of virulent viruses. Collectively, these findings have unveiled a rich diversity of novel viruses and their ecological functions within oil reservoirs. This study provides a comprehensive understanding of the role of viral communities in the biogeochemical cycle of the deep biosphere. It also facilitates microbial applications aimed at addressing challenges related to fossil-fuel production and carbon emissions in the petroleum industry.

## Introduction

The global dependence on fossil-fuel energy and the escalating levels of atmospheric carbon dioxide presents a pressing challenge to humanity^1^. In the realm of subsurface energy and carbon reservoirs, oil reservoirs have long harbored diverse microorganisms, a subject of growing interest since the 1930s^2^. Recent research has intensified the focus on understanding the presence and role of microorganisms in these deep subsurface environments^3–5^, driven by their dual impact on energy production and carbon emissions.

Despite extreme environmental conditions, oil reservoirs still offer a variety of niches to support various microorganisms^6^, including hydrocarbon oxidizing bacteria, fermentative bacteria, sulfate reducing microorganisms (SRMs), nitrate reducing microorganisms, and methanogens^7–11^. Microorganisms in oil reservoirs have demonstrated the ability to adapt to these extreme conditions and contribute to important ecological processes, such as the formation and transformation of crude oil, the geochemical cycle, and evolution of life^12,13^. On one hand, these microorganisms play a substantial role in the generation of ’heavy oil’^12^, leading to the deterioration of the world’s oil resources^3^ and significant carbon emissions during its recovery. On the other hand, microbial H_2_S production by sulfate reducing microorganisms (SRMs) is a prevalent metabolic process within the subsurface oil reservoirs^14^, which can lead to the corrosion of metal equipment and infrastructure, souring of oil, and risk of health, thus results in significant economic and ecological costs^14^. In addition, depleted hydrocarbon reservoirs represent the second-largest potential for CO_2_ storage^15^ and many hydrocarbon reservoirs have undergone CO_2_ injection as part of enhanced oil recovery (CO_2_-EOR) initiatives^16,17^. However, it’s worth noting that while microbial methanogenesis, another prevalent subsurface process, is considered a promising approach for bio-converting residual oil into CH_4_ in the petroleum industry, it also acts as a significant CO_2_ sink, potentially heightening the risk of gas loss during CO_2_ storage^18^.

While the importance of microorganisms within oil reservoirs has become increasingly recognized, the mechanisms governing the assembly and regulation of microbial communities in these deep subsurface environments remain unclear. Recent studies have shown that microbial communities in oil reservoirs undergo fluctuations during microbial enhanced oil recovery, and the reason for this phenomenon remain mysterious to date^19^. Additionally, variation partitioning of beta-diversity in oil reservoir reviewed that chemical properties and physical conditions explained a substantially larger fraction of variation in microbial beta-diversity. However, more than 70% of the community variation could not been explained^20^. Viruses are directly associated with host physiology and mortality and further influence microbial community dynamics in aquatic environments^21^. Therefore, viruses in the unique environment of oil reservoir may be one of the major contributors to the dynamic changes observed in the microbial community within these reservoirs.

Viruses are the most abundant biological entities on the planet and are found in nearly all environments^22^, even in extreme environments such as Antarctic soils^23,24^, thawing permafrost^25^, cryoconite hole^26,27^ and desert^28^. They play pivotal roles in natural ecosystems by interacting with microbial hosts and exerting significant influence over global biogeochemical cycles, such as global cycling of nutrients, energy flow, and food web dynamics^29^. Viruses can function as predators, regulating microbial abundance while releasing organic matter and inorganic nutrients through cell lysis^30^. Furthermore, viruses actively regulate and rebuild host metabolism by deploying auxiliary metabolic genes (AMGs) during infection. Recent studies have uncovered the presence of a substantial number of viruses within the oil reservoir environment^31,32^. A meta-analysis of eight production wells indicated that the abundance of viruses is approximately 3 × 10^8^ mL^−1^. Viral abundance of production wells during water flooding is higher than that in production wells during microbial flooding^31^. In addition, previous study uncovered that viruses are widespread in hydraulically fractured wells, rivalling the viral abundance observed in peatland soil ecosystems^33^. Repeated encounters with viruses and the shifting range of viral hosts occur across temporally and geographically distinct shale formations. Laboratory experiments showed that prophage-induced dominant microorganism lysis releases intracellular metabolites that can sustain key fermentative metabolisms^33^, supporting the persistence of microorganisms in this ecosystem. These findings reveal the potential significance of viruses in shaping microbial communities and reprograming microbial metabolisms within the oil reservoir ecosystem.

In addition, oil reservoirs harbor a broad diversity of uncultured microorganisms and novel metabolic pathways. Zhou et al. suggested that the archaeon ‘*Candidatus Methanoliparum*’ alone has the ability to degrade various large hydrocarbons into methane^7^. Meckenstock et al. discovered complex microbial communities that inhabit small water droplets within the oil phase, and it is widely believed that microbial degradation takes place in this zone^34^. Diverse functional gene groups identified from Pseudomonas in oil phase were significantly differed from those in the corresponding water phases^32^. These means that oil reservoirs are a unique evolutionary environment, containing uncultured microorganisms and their co-evolved novel viruses. Therefore, the study of oil reservoir virus is conducive to revealing global virus diversity.

However, the study on viruses in oil reservoirs is still in its early stage, and a comprehensive understanding of these viral communities remains elusive. Notably, there has been a notable absence of genome-level viral data from oil reservoirs to date. To bridge this knowledge gap and gain insights into the diversity, virus-host interactions, and potential ecological roles of viruses within oil reservoirs, it is imperative to establish a virome catalog specific to these environments. This endeavor is of paramount importance, especially in elucidating the mechanisms behind microbial impacts on energy production and carbon emissions in deep subsurface oil reservoirs.

## Results

### 1. Overview of prokaryotic genome and viral genome catalogues in the oil reservoirs

To explore the diversity and ecological function of viruses inhabiting oil reservoirs, we collected 59 oil reservoir produced water samples spanning China and performed whole shotgun metagenomic sequencing. In order to compile comprehensive catalogs of both prokaryotic and viral genomes from these metagenomic datasets, we devised a specialized pipeline (see Supplementary Fig. 1). Employing this pipeline, we analyzed the 59 metagenomes newly generated in this study, in addition to 123 metagenomes obtained from the public databases, originated from oil reservoir samples collected from Europe, Asia, North America, and South America (Fig. 1a). Consequently, we successfully constructed exhaustive catalogs of both prokaryotic and viral genomes. The prokaryotic genome catalogue comprised 7,229 medium-to high-quality microbial metagenome-assembled genomes (MAGs), encompassing 6,686 bacterial and 543 archaeal MAGs, spanning across 72 bacterial and 9 archaeal phyla. The bacterial community was dominated by Proteobacteria (2,402 MAGs), while the archaeal community was predominantly represented by Halobacteriota (339 MAGs). The viral catalogue of oil reservoir comprised a total of 33,657 putative viral genomes, all exceeding 10 kb in size (Supplementary Fig. 2a). This catalog featured 6,218 viral genomes of medium-to high-quality (> 50% completeness). To assess the coverage of viral communities within the oil reservoir, we conducted additional sequencing of virus-like particles (VLPs) from an additional 8 samples collected from oil reservoirs. The result revealed that 77.32% of the VLP reads could be mapped to the viral catalogue, suggesting that the viral catalogue extensively covered the viral population within the oil reservoir environment^35^.

**Fig. 1.**
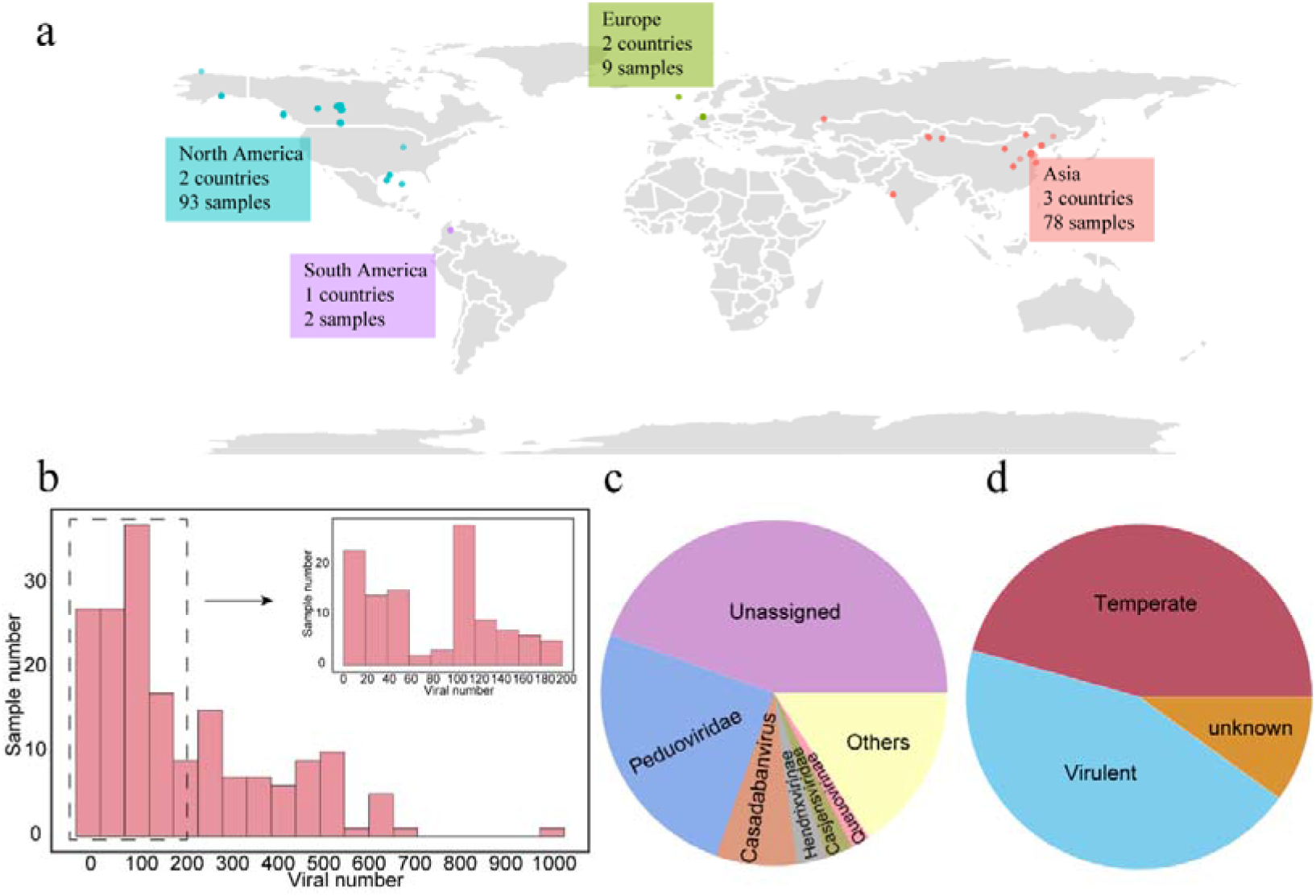
Overview of viruses in the oil reservoir ecosystem. a, Geographic distribution of collected oil reservoir samples. b, Histogram showing the distribution of vOTUs number across samples. c, Pie chart representing the relative proportion and taxonomy classification of vOTUs based on latest ICTV classification using PhaGCN2.0. d, The lifestyles of vOTUs within the oil reservoirs.

We clustered these 6,218 viral genomes into 3,886 viral Operational Taxonomic Units (vOTUs) with a sequence similarity threshold of 95%, a level that approximate species-level taxonomy^36^. The viral richness of all samples ranged from 1 to 1020 with an average of 206.11 (Fig. 1b). Rarefaction analysis showed that the number of detected vOTUs was saturated (Supplementary Fig. 2b), suggesting that our study provided a reasonably comprehensive sampling of viral communities within oil reservoirs. We assigned the vOTUs to taxonomic ranks based on the latest ICTV classification using PhaGCN2.0. Surprisingly, 44.52% of vOTUs could not be confidently annotated at the family level (Fig. 1c), in addition, only 5.20% vOTUs could be found in the IMG/VR v3 dataset, underscoring the presence of a substantial number of unidentified viruses within oil reservoirs. Among the annotated vOTUs, the predominant vOTUs were assigned to the Caudoviricetes class (formerly known as Caudovirales order, accounting for 43.46% of the total), including families such as Peduoviridae (25.06%), Casadabanvirus (7.54%), Hendrixvirinae (2.86%), and Casjensviridae (2.47%). Furthermore, we identified three core vOTUs that were present in more than 50% of samples. Notably, these core vOTUs represented a mere 0.08% of the total vOTUs. The majority of vOTUs (84.64%) were detected in less than 10% of the samples, highlighting the highly heterogeneous nature of viral communities within oil reservoirs. Additionally, our analysis predicted 1,722 and 1,774 vOTUs as virulent and temperate viruses, respectively, while the remaining vOTUs remained undetermined (Fig. 1d).

### 2. Biogeography of virial communities in oil reservoirs

To investigate how viral communities assembled within oil reservoirs, we compared the viral diversity and composition across various oil reservoir samples. We found significant disparities in viral diversity and composition associated with different geographic locations (Supplementary Fig. 3a, Fig. 2a, ANOSIM, R = 0.59, *P* = 0.001), This result suggested that the geographic location serves as the primary determinant of viral variation between samples on an intercontinental geographic scale. This discovery aligns with previous findings in studies of viral communities in cold seep and acid mine drainage environments^37,38^. Within the dataset employed in this study, the majority of samples from Canada and China oil reservoirs were dominated by Peduoviridae, while a few Chinese oil reservoir samples exhibited dominance by Casadabanvirus. The majority viruses of samples from USA oil reservoirs remained unclassified at the family level (Supplementary Fig. 3b).

**Fig. 2.**
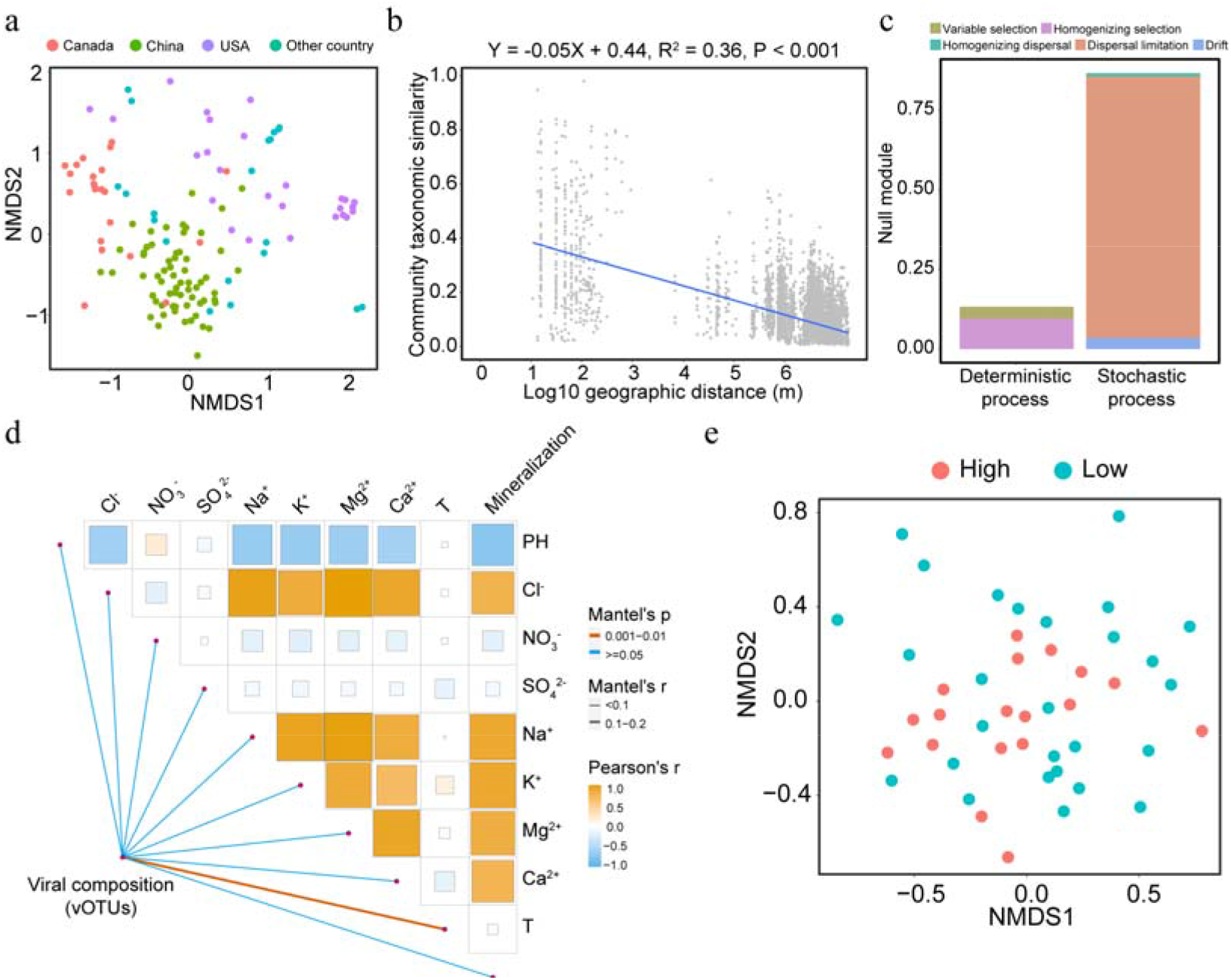
The distribution and assembly process of viral communities. a, Nonmetric multidimensional scaling (NMDS) of viral communities colored by sampling sites. b, Distance-decay relationships (DDRs) based on Bray-Curtis similarity (1 − dissimilarity of viral communities). The blue line denotes the least-squares linear regression across all spatial scales. Fitting equation, adjusted *R*^2^ values, and *P* values for DDR are presented in top. c, The proportion of viral community assembly process in oil reservoirs, including variable selection, homogenizing selection, homogenizing dispersal, dispersal limitations, and drift. d. Pairwise comparisons of the biotic and abiotic variables. The color and size of square in heatmap represents Pearson Correlation Coefficient (r). Edge width corresponds to the Mantel’s r statistic for the corresponding distance correlations, and edge color denotes the statistical significance. e. NMDS of viral communities from high and low temperature of oil reservoir.

Furthermore, we observed significant negative distance-decay relationships (DDRs) across all samples based on the Bray-Curtis similarities (1 − dissimilarity) of viral communities (slope = −0.05, *P* < 0.001) (Fig. 3b). To explore the mechanisms for viral community assembly within oil reservoirs, we performed a null model analysis. The result revealed that stochastic processes played a more substantial role than determinism in governing the viral community assembly in oil reservoirs (Fig. 2c). Moreover, we utilized 45 metagenomes newly sequenced in this work to examine how environmental factors drive viral community. Mantel tests indicated that the compositions of viral communities were significantly related to the temperature of oil reservoir (mantel statistic r = 0.12, *P* = 0.001) (Fig. 2d). NMDS analysis also illustrated significant dissimilarities in viral communities between high and low-temperature oil reservoirs (ANOSIM, R = 0.10, *P* = 0.018) (Fig. 2f).

**Fig. 3.**
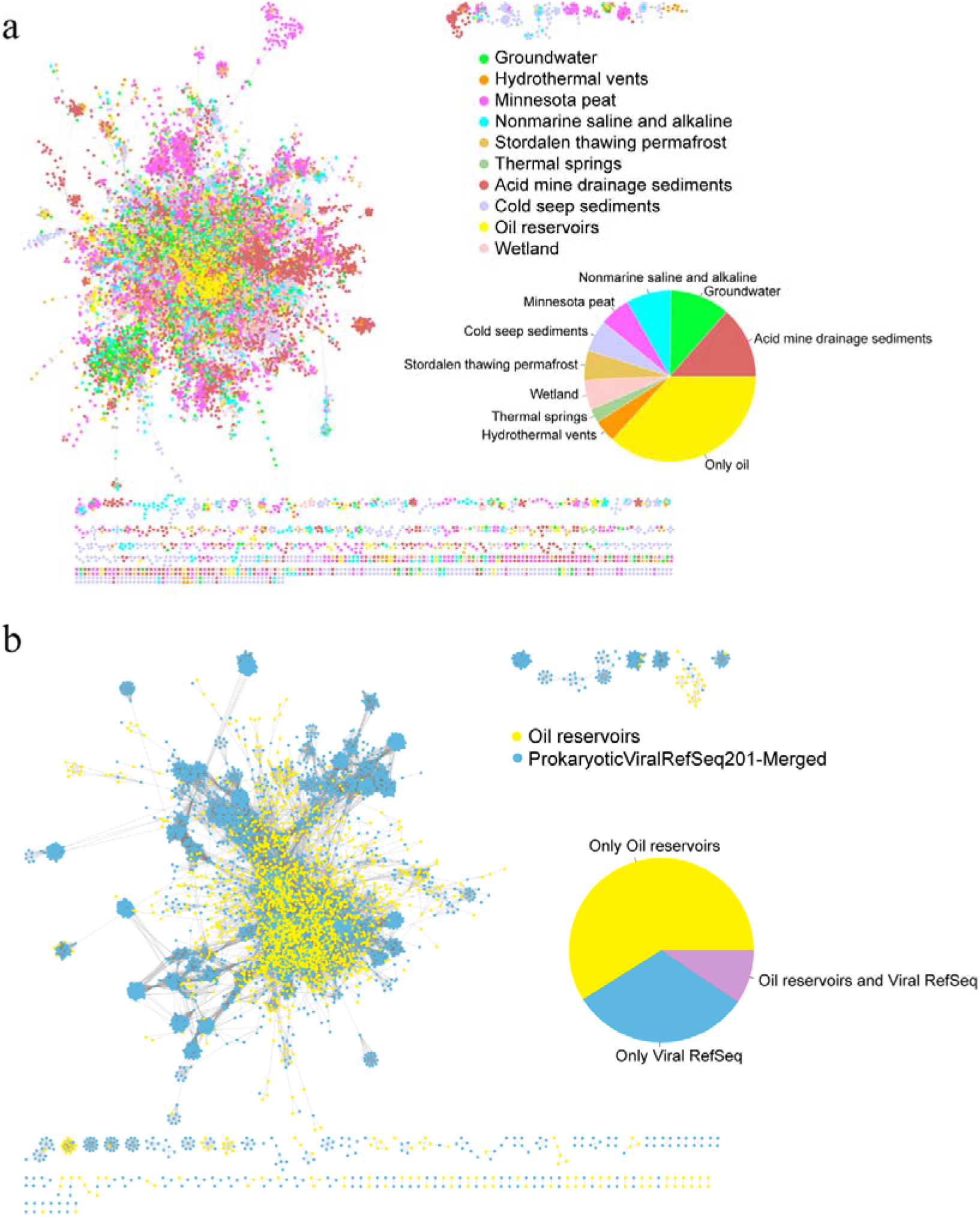
Taxonomic diversity of viruses in oil reservoirs. a, Gene-sharing network of viruses across various environments, including oil reservoirs, groundwater, cold seep sediments, thermal springs, wetland, acid mine drainage sediments, minnesota peat, non-marine saline and alkaline, hydrothermal vents, and stordalen thawing permafrost. The nodes in the network represent viruses, while the edges indicate similarity based on shared protein clusters. The color of the nodes represents the origin of viruses. Pie chart showing the relative proportion of shared viral clusters between oil reservoirs and other nine environmental virus datasets. b, Gene-sharing network of viruses from oil reservoirs and RefSeq prokaryotic viruses. Pie graph showing the relative proportion of shared viral clusters between oil reservoirs and RefSeq prokaryotic viruses.

### 3. Novel viral clusters in the oil reservoirs

To uncover novel viruses inhabiting oil reservoirs, we employed vConTACT2 to construct a gene-sharing network including vOTUs found in oil reservoirs and a wide variety of ecosystems, such as groundwater, sediment, and thermal springs, etc. In this weighted network, all vOTUs were grouped into 4,011 VCs (Fig. 3a). Among these VCs, 1,934 VCs were exclusively associated with a single ecosystem, while only two VCs were shared across all ecosystems. The limited overlap of viruses between different ecosystems demonstrated a high degree of habitat specificity among viruses. Within the subset of oil reservoir viruses, 2,942 out of 3,886 vOTUs were clustered into 805 VCs, with 265 VCs (32.92%) being unique to oil reservoirs. This finding suggested that the majority of oil reservoir viruses may be endemic to oil reservoirs. Furthermore, we found that acid mine drainage sediments and groundwater shared a higher number of VCs with oil reservoirs (Fig. 3a). Additionally, we also constructed the gene-sharing network based on the vOTUs found in oil reservoirs and those deposited in Viral RefSeq database. Our analysis showed that only a small percentage (n = 495, 14.66%) of vOTUs from oil reservoirs were clustered with taxonomically known genomes from Viral RefSeq (Fig. 3b). These results indicated that oil reservoirs possess a vast array of undetermined viruses.

### 4. Functional genes encoded by viromes in oil reservoirs

To unravel the functional roles virus within oil reservoirs, we clustered all 346,145 predicted protein-coding genes derived from oil reservoir viral genomes into 322,060 gene clusters. We found that 61.32% of viral gene clusters (accounting for 63.90% of total viral genes) lacked functional annotation against eggNOG database. The largest gene cluster derived from viromes in oil reservoirs predominantly encoded proteins associated with Replication, recombination and repair (L) functions. (Supplementary Fig. 4a).

Furthermore, we identified 105 putative auxiliary metabolic genes (AMGs) that might participate in the host metabolism and adaptation to the environment, mainly involved in Carbon utilization metabolism (n = 32), Energy metabolism (n = 25), miscellaneous metabolism (MISC, n = 34), and Transporters metabolism (n = 14). Among these AMGs, genes involved in Cobalamin biosynthesis (including *cobS* and *cobT*) were the most prevalent AMGs (Fig. 4, Supplementary Fig. 4b), which were found in 25 samples. Cobalamins are a class of structurally diverse cofactors containing cobalt^39^, essential for various biological functions, such as amino acid synthesis and carbon metabolism^40^. Importantly, microorganisms depend on cobalamins often relying on other species for cobalamins production, resulting in a network of cobalamin-dependent interactions^41^. These findings suggest that viruses could potentially influence microbial interactions by regulating the cobalamins production within their host organisms in oil reservoirs. In addition, we identified 9 AMGs from complete/high-quality vOTUs that were predicted to participate in energy metabolism (Supplementary Fig. 4c, d).

**Fig. 4.**
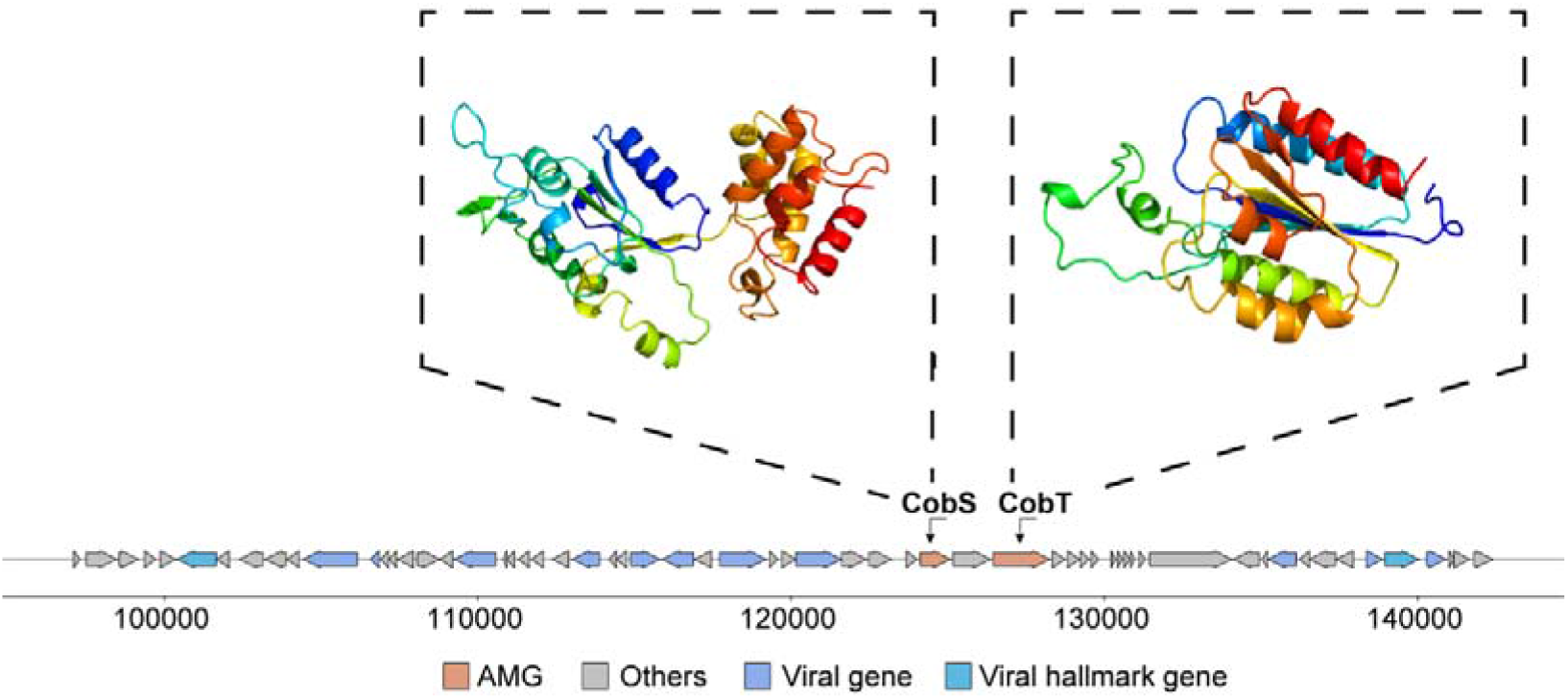
Genome map of representative *cobS* and *cobT* encoding viruses, AMGs are in orange, virus-like genes are in blue, viral hallmark genes are in dark blue, and non-virus-like or uncharacterized genes are in gray. Tertiary structures of selected AMGs based on structural modelling using Phyre2.

### 5. Close interactions between the virus and host in oil reservoirs

To further study the potential impacts of viruses on microbial ecology within oil reservoirs, we investigate in-situ virus-host interactions based on CRISPR spacers and tRNA sequences. A total of 7,197 putative virus-host pairs were predicted (1,411 based on CRISPR, 5,786 based on tRNA), in which 1,119 vOTUs (28.80% of the total vOTUs) were connected to 1,217 prokaryotic MAGs (host MAGs, accounting for 16.83% of the total MAGs). Most of the viruses with host links were not taxonomically assigned to known viruses at the family level (Supplementary Fig. 5b-d). The host MAGs were distributed in 3 archaeal phyla and 37 bacterial phyla (Fig. 5a, Supplementary Fig. 5a). The top five most frequently predicted bacterial phyla included Proteobacteria (785 MAGs), Actinobacteriota (69), Desulfobacterota (64), Chloroflexota (50), and Bacteroidota (49). The most frequently predicted archaeal phyla were Halobacteriota (10) and Methanobacteriota (8). In addition, we found 622 host MAGs with the potential for hydrocarbon degradation. These host MAGs have the ability to degrade alkanes and aromatics via aerobic or anaerobic degradation pathways. Two host MAGs were potential methanogenesis.

**Fig. 5.**
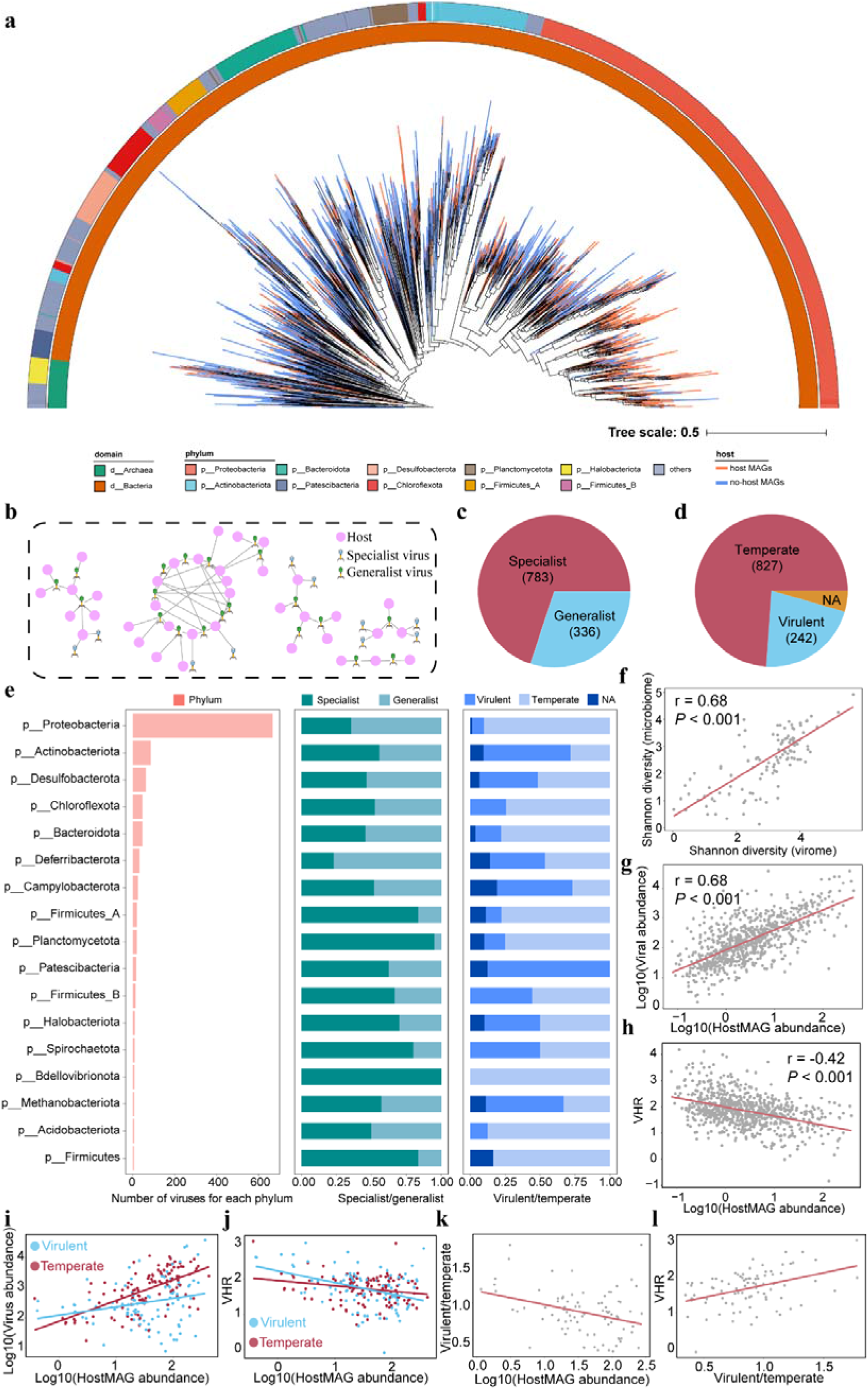
Linkages between Oil reservoirs viruses and host MAGs. a, Maximum-likelihood phylogenetic trees of MAGs detected in the oil reservoirs at the phylum level. The color of clades in the tree indicates whether MAGs are host MAGs (Orange represents host MAGs, blue represents non-host MAGs). The inner circle represents the domain, and the outer circle represents the phylum annotated by GTDB. The tree was constructed using PhyloPhlAn and visualized with iTOL. b, Schematic illustration depicting the relationships between host MAGs and their viruses. c, Pie chart showing the relative proportion of specialist and generalist viruses that can infect hosts. d, Pie chart showing the relative proportion of different types of lifestyle viruses that can infect hosts, including virulent, temperate, and undetermined (NA) viruses. e. Number of viruses, the ratio of specialist and generalist viruses, and the ratio of virulent and temperate viruses for each predicted host (at the phylum level). Seventeen phyla predicted to be infected with more than 5 viruses are shown in the figure. f, Comparisons of α-diversity (Shannon index) between the virome and the microbiome are presented. Regression lines are indicated in red. g, h, Correlation between the relative abundance of hosts and viruses (g) or VHR (h) in the oil reservoirs. Regression lines are indicated in red. i, j, Associations between the relative abundance of hosts and viruses (i) or VHR (j) (temperate and virulent viruses) in different samples. The red points represent temperate viruses, and the blue points represent virulent viruses, each point represent one sample. Regression lines are indicated in red and blue for temperate and virulent viruses, respectively. k, l, Correlation between the relative abundance ratio of virulent-to-temperate viruses and the relative abundance of hosts (k) or VHR (l). Regression lines are indicated in red.

Notably, the predicted host MAGs were also associated with several poorly characterized phyla, such as CPR (17), Sumerlaeota (4), Synergistota (4), Bipolaricaulota (3), Thermotogota (3), and one archaeal MAGs from the Thermoplasmatota phylum. Among these vOTUs with host links, 783 vOTUs (69.97% of the vOTUs with host links) was predicted to have a narrow host range, exclusively infecting unique hosts (termed specialist viruses). 336 (30.03%) vOTUs exhibited a broader host range, infecting more than one host (generalist viruses) (Fig. 5b, c).

Furthermore, the examination of viruses across their predicted hosts revealed significant differences in the proportion of specialist and generalist viruses within different host lineages. Lineage-specific virus-host interactions were enriched in Bdellovibrionota, Planctomycetota, and Firmicutes, while viruses infecting Deferribacterota and Proteobacteria tended to be generalist viruses (Fig. 5e).

Previous studies have suggested that viruses exert a significant influence on the structure of the microbial community through processes including both microbial lysis and integration as prophages. However, the relationships between viruses and microbes in the oil reservoir remain poorly understood. To investigate this relationship, we conducted a correlation analysis between viral and microbial profiles. Our analysis revealed a significant positive correlation of α-diversity (Shannon diversity) between viral community and prokaryotic community (r = 0.68; Fig. 5f). This result indicated that the structures of viral and prokaryotic communities are closely related within the oil reservoir environment. To further explore their associations, we examined the relationship between the relative abundance of viruses and host microbes. Our results unveiled a positive correlation between viral and microbial abundances (Fig. 5g). This finding suggested that viruses and their host microbes coexist in the oil reservoirs rather than being mutually exclusive. Furthermore, we noted a negative association between VHRs (Viral-to-Host Ratios) and microbial abundance within oil reservoir, that the hosts with higher relative abundances tended to exhibit lower VHRs (Fig. 5h). This phenomenon aligns with ‘piggybacking the winner’ (PtW) model, which posits that viruses exploit their hosts through lysogeny rather than killing them when host density is high. Temperate viruses protect their host cells from infection by closely related viruses via superinfection exclusion. Consequently, the contribution of temperate viruses to the host increases with high host abundance, resulting in a ‘more microbes, fewer viruses’ scenario.

To further clarify whether viruses affected their prokaryotic hosts in oil reservoirs through a ‘piggybacking the winner’ mechanism, we compared the abundances of temperate and virulent viruses with identified host links. Firstly, among the 2,493 temperate viruses, 827 were able to establish connections with their host MAGs in situ. In contrast, only 242 out of 3,180 virulent viruses were predicted to have host associations (Fig. 5d). Secondly, although both virulent and temperate viruses exhibited positive correlations in relative abundances with their hosts, the increase in the abundance of temperate viruses occurred at a higher ratio than that of virulent viruses (Fig. 5i), and the VHRs of temperate viruses exhibited a milder decline compared to those of virulent viruses (Fig. 5j). Thirdly, we found that the abundance of hosts exhibited a negative correlation with the ratio of virulent and temperate viruses (Fig. 5k), whereas the VHR displayed a positive correlation with the ratio of virulent and temperate viruses (Fig. 5l). Collectively, these findings suggested that the viruses affected their hosts mostly in the ‘piggybacking the winner’ manner within oil reservoirs.

Moreover, the proportion of temperate and virulent viruses varied among their hosts in oil reservoirs. While the majority of viruses predicted to infect hosts were temperate viruses, viruses predicted to infect Patescibacteria, Actinobacteriota, and Methanobacteriota were predominantly virulent. (Fig. 5e). In addition, we found that VHRs for most prokaryotic taxa exceeded one (Supplementary Fig. 6a-d), providing evidence of active viral genome replication within these lineages.

### 6. Sulfur metabolism mediated by viruses in oil reservoirs

In oil reservoirs, a prevalent metabolic pathway involves the microbial reduction of sulfate to generate hydrogen sulfide (H_2_S), a process primarily conducted by sulfur-reducing microorganisms (SRMs)^42,43^. H_2_S has significant implications for the degradation of infrastructure, reservoir souring, the operation cost of oil production, and the value of crude oil^44^. Recently studies reported that viruses infect sulfur-metabolizing microbes and reshaping sulfur metabolism within host cells by encoding auxiliary metabolic genes (AMGs)^45^. To investigate the potential impact of viruses on microbial sulfur metabolisms in oil reservoirs, we identified the MAGs with sulfur metabolic abilities and their associated viruses. A total of 484 host MAGs were predicted to possess sulfur metabolic abilities, including assimilatory sulfate reduction (ASR), thiosulfate oxidation metabolism (TSO), dissimilatory sulfate reduction (DSR), and sulfide oxidation (SO) (Fig. 6a, Supplementary Fig. 7c, see supplemental material for more information).

**Fig. 6.**
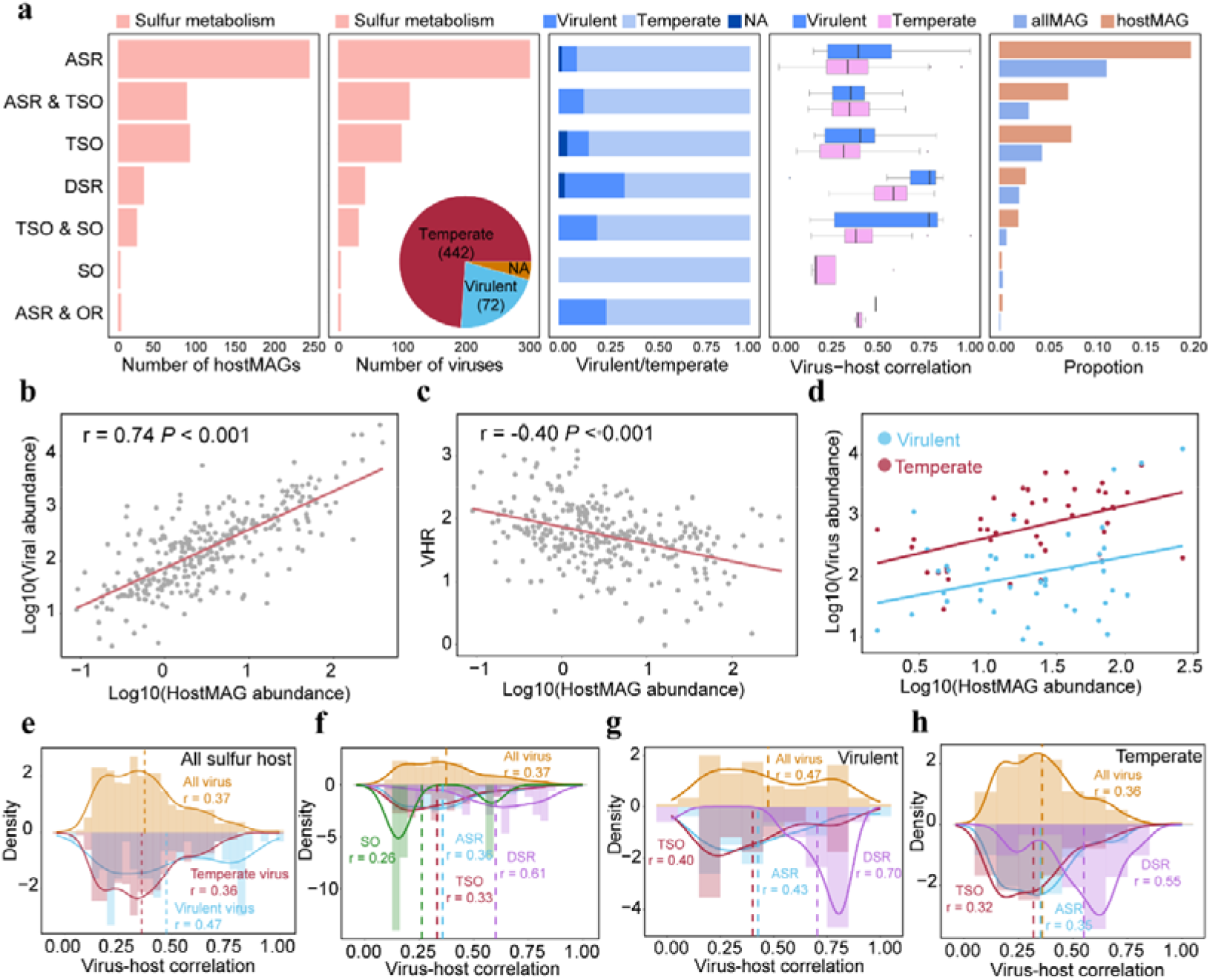
The impact of viruses on the sulfur cycle in oil reservoirs. a, Number of host MAGs, Number of viruses, the ratio of virulent and temperate viruses, spearman correlations between the relative abundances of viruses and predicted hosts, and the number proportion of MAGs in all MAGs and host MAGs for each sulfur metabolism ability function group. Pie graph showing the relative proportion of different lifestyle viruses that can infect the host, including virulent, temperate, and undetermined (NA) viruses. b, c, Correlation between the relative abundance of hosts with sulfur metabolism ability and the viruses (b) or VHR (c). Regression lines are indicated in red. d, Associations between the relative abundance of hosts with sulfur metabolism ability and the virus (temperate and virulent viruses) in different samples. Red points represent temperate viruses, blue points represent virulent viruses, every point represent a single sample. Regression lines are shown in red for temperate and blue for virulent viruses. e, Distribution of virus-host correlations with sulfur metabolism ability. Orange, blue, and red colors represent the distributions of all viruses, virulent, and temperate viruses, respectively. f, Comparison of distribution of virus-host correlations with different sulfur metabolism ability. Orange color represents the distributions of all viruses, while blue, green, purple, and red colors represent viruses infecting different function groups of sulfur metabolism. g, h, Distribution of virulent (g) or temperate (h) viruses and host correlations with sulfur metabolism ability. Orange color represents the distributions of all virulent (g) or temperate (h) viruses, while blue, purple, and red colors represent virulent (g) or temperate (h) viruses infecting different function groups of sulfur metabolism. Dashed lines show the average correlation in the distribution.

To assess the significance viruses in the context of sulfur metabolism within oil reservoirs, we quantified the relative proportion of MAGs with potential sulfur metabolism abilities within the entirety of MAGs and specifically among host MAGs. The result showed that the relative proportion of MAGs with potential sulfur metabolism abilities within host MAGs was higher compared to the overall MAGs population (Fig. 6a). Moreover, we found that related abundance of host MAGs with sulfur metabolism abilities and their associated viruses were positively correlated (Fig. 6b). These findings underscore the pivotal role played by viruses in the sulfur cycle of oil reservoirs.

To further unravel the impact of viruses on hosts with sulfur metabolism abilities, we explored the potential roles of temperate and virulent viruses in microbial sulfur metabolisms We found that most of the viruses infecting MAGs with sulfur metabolism abilities were temperate viruses (Fig. 6a) and the VHRs exhibited a decline corresponding to the increasing abundance of hosts with sulfur metabolism abilities (Fig. 6c). These results were consistent with the findings observed in the overall MAGs population. However, we noticed a parallel increase in the relative abundances of virulent and temperate viruses in tandem with the rise in host abundance (Fig. 6d). This difference in abundance increase rate between virulent and temperate viruses was not as pronounced as the trend observed in the entire host population (Fig. 5j, Fig. 6d).

In addition, virulent viruses displayed a significantly stronger correlation with their hosts engaged in sulfur metabolism compared to temperate viruses (Fig. 6e, Supplementary Fig. 7d-f). Collectively, these findings point to a relatively more substantial contribution of virulent phages to the regulatory dynamics of sulfur metabolic communities.

Intriguingly, we found that the proportion of virulent viruses predicted to infect hosts in DSR was higher than in the context of other sulfur metabolism processes (Fig. 6a). Hosts with DSR abilities also exhibited a relatively stronger correlation with their associated viruses compared to other hosts (Fig. 6f). This high correlation could be attributed to the higher prevalence of virulent viruses infecting these hosts, in contrast to other hosts, such as host with ASR ability, which are more frequently infected by temperate viruses (Fig. 6a). In addition, both virulent and temperate viruses infecting hosts with DSR ability showed a relatively stronger correlation with their hosts compared to other hosts (Fig. 6g, Fig. 6h). These findings collectively suggested that the viruses primarily regulated DSR function within oil reservoirs, possibly in a top-down regulatory manner^46^, where the growth and abundance of SRMs are primarily regulated by the predation of virulent viruses.

### 7. Evidence of viruses mediated sulfide production in microcosms

To further investigate the influence of viruses on sulfate reduction, we employed oil reservoir production water from the Huabei Oilfields as the initial inoculum and set up two distinct sets of microcosms, each characterized by varying initial concentrations of virus-like particles (VLPs) (Supplementary Fig. 12). Specifically, the SV microcosms featured a higher initial count of VLPs in comparison to the SM microcosms. Over the course of 570 days, we monitored H_2_S production and the viral communities were analyzed at two time points (160 days and 570 days) after the start of incubation (i.e., T1 and T2). Detailed information of putative viral contigs and host MAGs can be found in Supplemental File.

Throughout the incubation period, we observed a significant decrease in sulfide production in SV microcosms featuring a high initial count of VLPs (Fig. 7a). Additionally, at the T1 time point, the diversity of both overall microbial community and host-MAG subcommunity was significantly higher in SV microcosms compared to SM microcosms (Supplementary Fig. 8c).

**Fig. 7.**
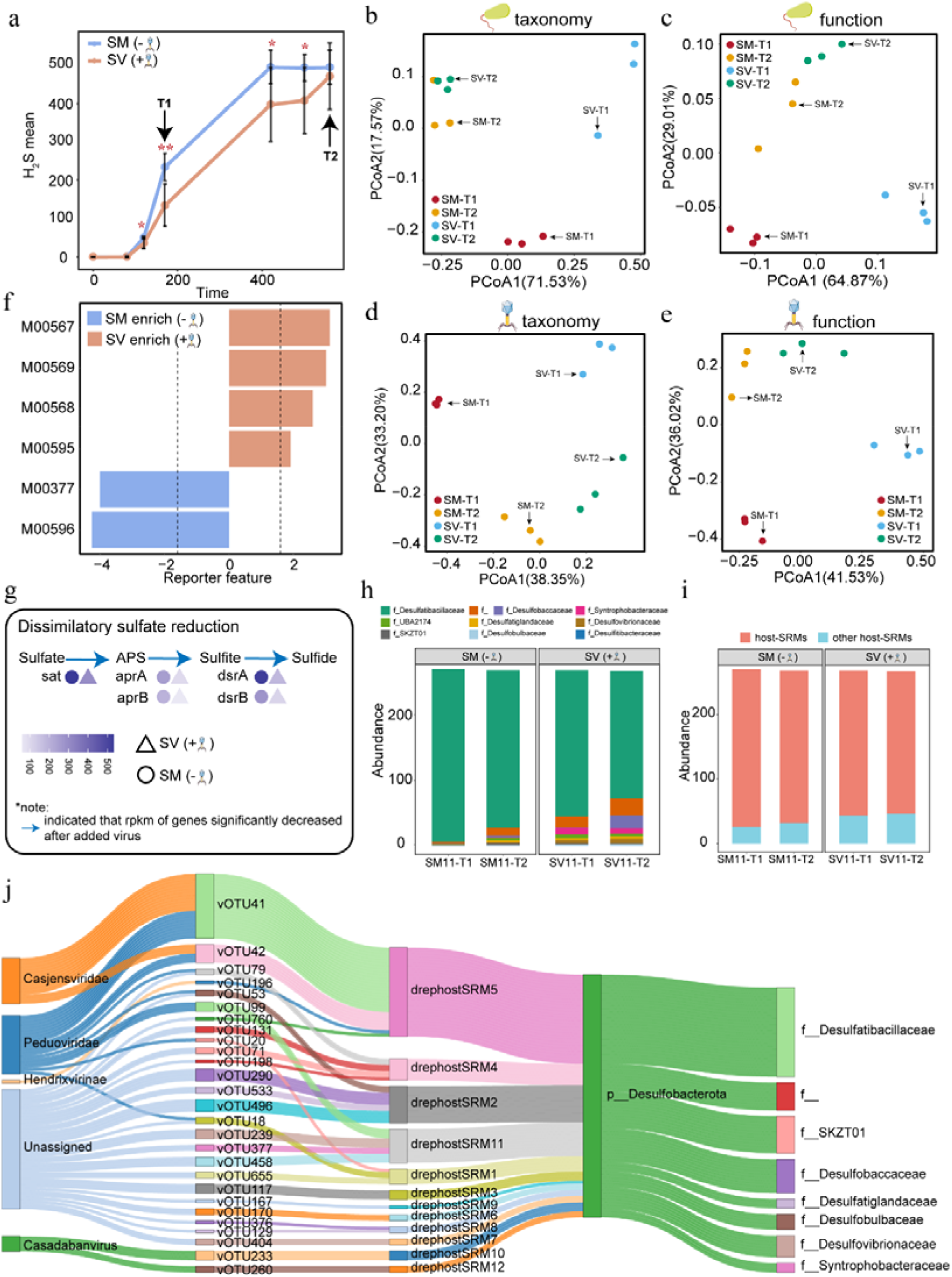
Viral impact on dissimilatory sulfate reduction. a, Sulfide production in microcosms. The blue asterisks indicate the significance level of the sulfide production comparing the SM microcosms with the SV microcosms at the corresponding time point (* *P* < 0.05, ** *P* < 0.01, *** *P* < 0.001). b, c, Changes in microbial community composition (b) and function (c) at T1, and T2 sampling time points. d, e, Changes in viral community composition (d) and function (e) at T1, and T2 sampling time points. f. Reporter score evaluated the enrichment of modules between the two types microcosms. g, Comparison of the abundance of dissimilatory sulfate reduction related genes in the SM and SV microcosms at T1 sampling time. The gradient dark blue circle represents the average RPKM value of the genes in the SM microcosms. The gradient dark blue triangle represents the average RPKM value of the genes in the SV microcosms. h, The abundance variation of all-SRMs in microcosms at the family level. i. The abundance proportion between host-SRMs and other host-SRMs in microcosms. j. Predicted virus-host links between virus and host SRMs in microcosms. The right two panels represent host taxonomy colored by phylum and family, and the left two panel show viral taxonomy colored by family, and viral clusters, connecting lines show associations between host SRMs and viruses.

To examine the potential influences of the initial counts of VLPs on the composition and function of both microbial and viral communities, we conducted Principal Coordinates Analysis (PCoA) to evaluate the degree of differentiation between the two sets of microcosms. The PCoA analysis revealed distinct dissimilarity patterns in both the composition and function of microbial community across the two microcosms (ANOSIM, R_composition_ = 0.85, *P*_composition_ = 0.001, R_function_ = 0.91, *P*_function_ = 0.001) (Fig. 7b, Fig. 7c). Similar to microbial community, the composition and function of viral communities followed different phases (ANOSIM, R_composition_ = 0.95, *P*_composition_ = 0.001, R_function_ = 0.96, *P*_function_ = 0.001) (Fig. 7d, Fig. 7e). As the incubation duration progressed, the differences in diversity and composition of microbial and viral communities between the two sets of microcosms diminishes (Fig. 7b-e, Supplementary Fig. 8d). Moreover, we compared the functional characteristics related to energy metabolism and hydrocarbon degradation within the SM and SV microcosms. Notably, the M00569 module, associated with DSR function, exhibited enrichment in the SM microcosms at T1 (Fig. 7f). Additionally, the abundance of genes involved in DSR was significantly lower in the SV microcosms compared to the SM microcosms at T1 (Fig. 7g). These results indicated that the high count of VLPs significantly reduced H_2_S production by diminishing the abundance of genes involved in DSR in the microcosms.

To gain insight into the potential influences of the initial counts of VLPs on the SRM community, we compared the abundance and composition of SRM across the different sets of microcosms. In total, we identified 89 SRMs (referred to as all-SRMs) including 44 SRMs associated with identified viruses (referred to as host-SRMs), accounting for 11.68% and 5.77% of total MAGs, respectively (Supplementary Fig. 9). In the SV microcosms, the SRM subcommunity displayed a higher diversity than the SM microcosms, which was similar with the findings observed within the entirety microbial community (Fig. 7h, Supplementary Fig. 10a, b). This result could be attributed to the decreased abundance of the dominant SRMs (drephostSRM5, *Desulfatibacillaceae*) in the community, which in turn led to an increase in the abundance of rare SRMs, such as drephostSRM3 (*Desulfatiglandaceae*), drephostSRM4 (*Unclassified Adiutricales*), and drephostSRM10 (*Syntrophobacteraceae*) (Fig. 7j).

Collectively, the results suggested that the high initial count of VLPs restructured the sulfate reduction microbial subcommunity in the SV microcosms. In addition, we compared the total abundance of SRM within the overall microbial community in different sets of microcosms. We found that the total abundances of SRM in SM were significantly higher than in SV microcosms at T1 (*P* = 0.024). Intriguingly, we also found that the majority of the SRMs could be associated with identified viruses (Fig. 7i), suggesting that viral infection targeting SRMs was a primary reason for the reduction of H_2_S production. The high initial VLP count primarily attenuated H_2_S production by regulating the composition of SRMs.

## Discussion

In this study, we conducted large-scale data mining of metagenomes to construct a catalog of viromes form oil reservoirs. This extensive resource contains a wide range of viral genomic diversity that is unique to oil reservoirs. It encompasses diverse and previously uncharacterized viral groups. In addition, we preliminary verify the potential impact of viruses on the sulfate reduction microbial subcommunity. These findings will contribute to a better understanding of the ecological role of viruses in deep biosphere, such as regulating microbial mortality, structuring microbial community, impacting biogeochemical cycling, and mediating microbial evolution (Supplementary Fig. 11).

Recent metagenomic and virome surveys have uncovered a remarkable range of viruses in both aquatic and terrestrial environments, significantly enhancing our understanding of virome diversity^37,47–49^. However, our knowledge of viral communities in oil reservoir is still limited. Most of the identified viruses from oil reservoir were novel genomes that not previously characterized. Moreover, the taxonomic annotation ratio of oil reservoir vOTUs was lower than that of seawater-derived (26.16%) and sediment-derived (15.18%) vOTUs^50^. This low annotation proportion can primarily be attributed to the absence of complete genomes from viral isolates in oil reservoirs and associated environments. Collectively, our findings demonstrate the uniqueness of viruses in oil reservoirs, emphasizing the limited extent of our knowledge regarding viral diversity in these environments.

Comparisons between viruses from oil reservoirs and other ecosystems can provide valuable insights into oil reservoir viral ecology. Gene-sharing network analysis showed that viruses from oil reservoirs and other ecosystems formed separate cohesive clusters, suggesting that viruses may possess unique metabolism gene to adapt to oil reservoir environments. Oil reservoirs are a relatively independent and stable ecosystem that have been isolated over millions of years, with unique microorganisms inherent in them^51^. The co-evolution of viruses and microorganisms may lead to distinct nature of oil reservoir viromes compared to those reported in other environments. This underscoring the scarcity of research and datasets on viromes in oil reservoirs. In addition, we found that groundwater and acid mine drainage sediments shared more VCs with oil reservoirs. Groundwater is also part of the deep biosphere. Oil reservoirs and groundwater are closely interrelated and inseparable, characterized by the profound interplay between geomicrobiology within petroleum-contaminated aquifers and the chemistry of groundwater^52^. Therefore, viruses from oil reservoirs and groundwater share more common genes. As for acid mine drainage sediments, despite the significant environmental differences between oil reservoirs and acid mine drainage sediments, it is speculated that viruses in these two environments may either have similar origins or possess adaptation strategies to acidic/alkaline environments.

The proportion of temperate viruses in oil reservoirs is higher compared to other ecosystems, such as seawater, sediment, and soil^47,53^. Several studies have found that the temperate strategy is advantageous in oligotrophic conditions^54^. Temperate viruses provide benefits to their hosts though various mechanisms, such as genome engineering and the regulation of gene expression and function^55^. Moreover, temperate viruses can enhance the fitness and competitiveness of hosts, helping them thrive in oligotrophic environment by altering cell physiology and introducing novel functions^55^. Hence, in harsh environments, the temperate strategy may serve as an effective adaptive mechanism for viruses, enabling their long-term coexistence with hosts in oil reservoirs.

The associations between viral and host abundance have been described by the Kill-the-Winner (KtW) and Piggyback-the-Winner (PW) theories^54,56–58^. Density- and frequency-dependent lytic KtW models predict that a high bacterial abundance is associated with a high rate of lytic infections, leading to an increased VHRs^58^. On the other hand, PW theory suggests that temperate viruses can protect their host cells from closely related viruses via superinfection exclusion, thus at high host densities, rather than killing their hosts, viruses might switch their lifestyle from virulent to temperate life cycle and replicate integrated into their host genomes, resulting in VHRs decreased^54,58^. This trend has been observed in various ecosystems, from soil to freshwater to human lungs^59,60^. In oil reservoirs, the associations between viral and host abundance also support the PW theory. The ratio between virulent and temperate viruses decreases with increasing host density (Fig. 5k), suggesting that temperate viruses are a more successful strategy for viral replication at high host densities in oil reservoirs.

SRMs are an important microbial group in oil reservoirs^42^. Our association analysis showed that hosts with dissimilatory sulfate reduction abilities have the highest correlation with viruses compared to hosts involved in other sulfur metabolism. This high correlation can be explained by the higher proportion of virulent viruses that infect hosts with dissimilatory sulfate reduction abilities compared to other sulfur metabolism hosts. Thus, virulent viruses play an important role in the dissimilatory sulfate reduction of the deep biosphere. To further validate this conclusion, we conducted a microcosmic experiment and discovered that viruses decrease sulfide production by reducing the abundance of genes involved in dissimilatory sulfate reduction. Additionally, we observed that the sulfate reduction microbial subcommunity following the ‘kill the winner’ model. In the microcosms with a high number of viruses, the relative abundance of dominant SRMs decreased due to viral infection. This release of niche space fostered the growth of other SRMs with a lower abundance. Therefore, in this way, viruses regulate the diversity and structure of the sulfate reduction microbial subcommunity. In summary, viruses not only inhibit the growth of SRMs, but also shape the structure of the sulfate reduction microbial subcommunity. Hydrogen sulfide (H_2_S) produced by SRMs not only causes souring of oil reservoirs, but also influences the cost of oil production and the value of crude oil^44^. In further oil exploration and production, it may be possible to introduce viruses for the prevention and treatment of souring in oil reservoirs.

Biodegraded oils constitute the majority of the global petroleum reserves^12^. The quality and value of the oil greatly depend on the extent of biodegradation and the presence of metabolic byproducts resulting from microbial activity, such as sulfur compounds and organic acids^3^. Furthermore, microbial enhanced oil recovery (MEOR) emerges as a cost-effective approach to extract remaining oil reserves^61^. Therefore, comprehending the microbial processes occurring in deep oil reservoirs is crucial for the advancement of MEOR techniques. Our findings indicate that over half of host MAGs exhibit the potential to degrade hydrocarbons, suggesting that viruses might also play a significant role in hydrocarbon degradation within oil reservoirs.

In conclusion, this study presents a viral catalogue of oil reservoirs, unveiling a rich diversity of novel viruses in oil reservoirs. Furthermore, our study elucidates a wide array of host-virus interactions and provides evidence for the substantial impact of viruses on the microbial sulfate reduction within oil reservoirs. These results sheds light on the ecological roles of viruses and their hosts in oil reservoirs. Our work also facilitates microbial applications aimed at addressing challenges related to fossil-fuel production and carbon emissions in the petroleum industry.

## Methods

### Collection of metagenomic data sets for oil reservoirs

A total of 59 oil reservoir production water samples were collected from five provinces across China. All samples were collected from the wellheads of each production well, where the oil and water mixture fluid were pumped out. Mixture fluid from wellheads was collected directly into clean and sterilized 5 L sampling bottles till the bottles were filled up to exclude oxygen. All samples were kept in an icebox and transported to the laboratory immediately and stored at 4 °C for DNA extraction. For better separation of oil and water from the mixture fluid, all the bottles filled with production mixtures were stood with gravitational precipitation for 12h at 4°C. Subsequently, five hundred milliliters of water phases were collected for each sample for total microbial genomic DNA extraction. Microbial cells were obtained after filtering through 0.22-μm-pore-size polycarbonate membranes (45 mm diameter; Millipore, Bedford, MA, United States). The polycarbonate membranes with the collected microbial cells were cut into small pieces by using the sterile scissor, and placed into the sterile centrifuge tubes for DNA extraction (FastDNA^®^ SPIN Kit for Soil, MP Biomedicals, USA), following the manufacturer’s instructions. Extracted DNA was used for library preparation with NEB Next^®^ Ultra II^TM^ DNA Library Prep Kit for Illumina^®^ (New England Biolabs, USA) and sequenced on Illumina NovaSeq 6000 platform (150 bp, paired-end reads). This generated totally ∼5 Tb metagenomic raw reads data. In addition, 123 publicly available oil reservoirs metagenomes datasets were downloaded from NCBI Sequence Read Archive (SRA) database in Nov 2021 (Supplementary Table S1).

### Collection of virus-like particles (VLPs) and extraction of viral DNA

To estimate whether the metagenome derived viral genome catalogue could cover the viral communities in oil reservoirs, we collected virus-like particles (VLPs) and sequenced viral DNA of additional 8 samples from oil reservoirs. Briefly, after obtained filtrates from water phase, the filtrates were further used for the extraction of VLPs. To obtain VLPs, the filtrates were firstly filtered through tangential flow filtration equipment (TFF) with a 100 kDa tangential flow membrane package (Sartorius, VIVAFLOW 50 100,000 MWCO, Germany), and then continuously concentrated until ∼1 mL solution using 100 kDa centrifugal filter units (Amicon^®^ Ultra-15, Ultracel-100K, Millipore, Germany). 720 μL concentrated solution was treated with 1000U/mL DNase I (37°C, 2h) (Roche, China) before viral DNA extraction. Total viral DNA was extracted using phenol-choroform-isopentanol method. Then, extracted total viral DNA was used to construct sequencing libraries and sequencing. Firstly, extracted total viral DNA was amplified using REPLI-g Cell WGA & WTA Kit (Qiagen, Germany), the products were then used to construct sequencing libraries using the NEB Next^®^ Ultra II™ DNA Library Prep Kit for Illumina^®^ (New England Biolabs, USA) following the manufacturer’s recommendations.

### Processing of metagenomic sequence data and generation of prokaryotic metagenome-assembled genomes

The metagenomic raw reads were examined using FastQC v0.11.9 (http://www.bioinformatics.babraham.ac.uk/projects/fastqc/), low-quality sequences, primers, and adaptors were trimmed using the Trimmomatic v0.39^62^ (parameters: LEADING:2 TRAILING:2 SLIDINGWINDOW:4:20 MINLEN:50). The trimmed reads were independently assembled using MEGAHIT v1.2.9^63^ (parameters: --presets metasensitive) and/or SPAdes v3.15.4^64^ (parameters: -meta, -k 21,33,55,77,99,127). For each assembly, contigs were binned using the binning module (parameters: --metabat2 --maxbin2 --concoct) and consolidated into a final bin set using the Bin_refinement module (parameters: -c 50 -x 10) within metaWRAP v1.2.1^65^. All the produced bin sets were aggregated and de-replicated at 95% average nucleotide identity (ANI) using dRep v3.2.2^66^ (parameters: -comp 50 -con 10 -nc 0.30 -pa 0.9 -sa 0.95). The completeness and contamination of MAGs were assessed using lineage-specific module within CheckM v1.1.3^67^ with default parameters.

The taxonomy of each MAG was assigned using GTDB-Tk v1.5.0^68^ based on the Genome Taxonomy Database (GTDB, http://gtdb.ecogenomic.org) taxonomy (release202). The maximum-likelihood phylogenetic trees of MAGs were constructed based on a concatenated dataset of 400 universally conserved marker proteins using PhyloPhlAn v3.0.64^69^ and visualized using iTOL v5^70^. RPKM (Reads per kilobase per million mapped reads) values were used to represent the relative abundances of MAGs. The RPKM values of the MAGs were calculated using CoverM v0.6.1 (https://github.com/wwood/CoverM) (parameters: coverm genome --min-read-percent-identity 0.95, --min-read-aligned-percent 0.75, --contig-end-exclusion 0 and -m rpkm).

### Functional annotation of MAGs and phylogenetic analysis of DsrAB protein

Open reading frames (ORFs) of these MAGs were predicted with Prodigal v2.6.3^71^ (parameters: -m meta). The predicted ORFs were annotated using eggNOG-mapper v2.0.1^72^ and the eggNOG Orthologous Groups database (version 5.0)^73^ in diamond mode. Annotated KO numbers were used for inferring the pathway encoded in each MAG. MAGs that encode complete pathway for hydrocarbon degradation, methane metabolism, and sulfur metabolism were utilized for further analyzed. In addition, to identify genes involved in the anaerobic degradation of hydrocarbons, we constructed a gene database involved in anaerobic degradation of hydrocarbons. Sequences from hydrocarbon-degrading enzymes in the AnHyDeg database (https://github.com/AnaerobesRock/AnHyDeg) and recently published protein sequences of alkyl-succinate synthases^8,74^. All proteins from MAGs were aligned against this database using BLASTp (-k 1 -e 1e-20 --id 30 --query-cover 70 --more-sensitive)

For phylogenetic analysis of DsrAB sequences, both DsrAB sequences obtained from MAGs and refDsrAB sequences reported previously^14,75^ were utilized for the phylogenetic analysis (Supplementary Fig. 7a, b, Supplementary Fig. 9), which could help to distinguish reductive and oxidative type DsrAB. The DsrAB sequences were aligned using MUSCLE v3.8^76^ with default parameters. The alignments were then filtered using TrimAL v1.4^77^ (parameters: -cons 50). The concatenated DsrAB tree was constructed using RAxML^78^ (parameters: -f a -m PROTGAMMAIJTT -p 12345 -x 12345 -N 100). The Newick files with the best tree topology were visualized using iTOL v5^70^.

### Viral contigs identification, dereplication, virus operational taxonomic unit (vOTUs) clustering, and calculating abundances

Viral contigs were recovered from assembled contigs using VirSorter v2.1^79^ and DeepVirFinder v1.0^80^. Only viral contigs ≥ 10 kb were retained, based on the following criteria: (1) Identified viral contigs only by VirSorter v2.1^79^ (parameters: --exclude-lt2gene); (2) Identified viral contigs only by DeepVirFinder v1.0^80^ (parameters: score ≥ 0.9 and p < 0.05); (3) Both identified by VirSorter v2.1^79^ and DeepVirFinder v1.0^80^. The identified viral contigs from each sample were clustered into virus operational taxonomic unit (vOTUs) using the parameters 95% average nucleotide identity (ANI) and 85% alignment fraction of the smallest scaffolds based on the scripts (https://bitbucket.org/berkeleylab/checkv/src/master/) provided in CheckV v0.8.1^81^. vOTUs were then detected provirus boundaries and removed host contamination using CheckV v0.8.1^81^. RPKM values were used to represent the relative abundances of vOTUs. The RPKM values of vOTUs were counted using CoverM v0.6.1 (parameters: coverm contig --min-read-percent-identity 0.95, --min-read-aligned-percent 0.75, --contig-end-exclusion 0 and -m rpkm). Viral lifestyle was predicted by both VIBRANT v1.2.1^82^ and CheckV v0.8.1^81^, while the remaining vOTUs with at least 90% completeness that display no prophage signals or lysogeny-specific genes were considered as potential virulent viruses. The detailed description of viral community, and statistical analysis performed in this work can be found in the Supplemental Materials and methods section.

### Viral taxonomic assignments, viral function annotation and identification of auxiliary metabolic genes(vAMGs)

Open reading frames (ORFs) of vOTUs were predicted with Prodigal v2.6.3 (-p meta -g 11 -f gff). To understand the taxonomy of vOTUs, we used PhaGCN2.0^83^ with the latest ICTV classification to explore the taxonomic affiliation of vOTUs at the family level. Additionally, to understand the function of vOTUs, the predicted viral proteins were first merged and dereplicated using CD-HIT v4.7^84^ (parameters: -c 0.90 -s 0.8 -n 5 -g 1 -d 0). The dereplicated viral proteins were assigned to the eggNOG Orthologous Groups database (version 5.0) using eggNOG-mapper v2.0.1 (-m diamond) to identify COG functional classifications. Moreover, we used DRAM-v v1.3.5^85^ to recover putative AMGs from vOTUs (See supplemental method for more information).

### Network analysis

Protein-sharing network analysis of vOTUs was performed by vConTACT v.2.0^86^. Briefly, all vOTUs of the oil reservoir were compared to vOTUs (≥ 10 kb) from other ecosystems in published paper data: (1) Wetland sediment (n = 1,075)^87^; (2) Stordalen thawing permafrost (n = 1,682)^25^; (3) Acid mine drainage sediments (n = 5,184)^38^; (4) Cold seeps (n = 2,490)^37^, (5) Minnesota peat (n = 3,566)^88^; and from other ecosystems in IMG/VR v3^89^: (6) Hydrothermal vents (n = 531); (7) Groundwater (n = 1,372); (8) Non-marine Saline and Alkaline (n = 1,756); (9) Thermal springs (n = 268). For each vOTUs, ORFs were called using Prodigal v2.6.3^71^, and the predicted protein sequences were used as input for vConTACT v2.0^86^. The protein sequences of the vOTUs were grouped into protein clusters (PCs) using vConTACT v2.0^86^ (parameters: --rel-mode Diamond --pcs-mode MCL --vcs-mode ClusterONE). The degree of similarity between the vOTUs was calculated based on the number of shared PCs. The networks were visualized by Cytoscape v3.5.1^90^ (http://cytoscape.org) using an apply preferred layout model. Additionaly, all vOTUs of the oil reservoir were also compared to NCBI Prokaryotic Viral RefSeq v201 database using vConTACT v2.0^86^.

### Virus-host prediction

Two different in silico methods were used to predict virus-host interactions. (1) tRNA match. ARAGORN v 1.2.38^91^ was used to identify tRNAs from sequences of vOTUs (parameters: -t). Identified tRNAs were compared to metagenomic contigs using fuzznuc^92^ from the EMBOSS:6.6.0.0 package with no mismatches allowed. (2) CRISPR spacer match. CRISPR spacers were recovered from metagenomic contigs using metaCRT (modified from CRT1.2)^38,93^ with default parameter. Extracted spacers were compared to vOTUs using fuzznuc^92^ from the EMBOSS:6.6.0.0 package with no mismatches allowed^94^.

### Microcosm experiments setup

To further study the viral impact on sulfate reduction, microcosm experiments were performed. We used the production water samples collected from the Huabei Oilfields (Block B51-11, 38 °C) as seed banks, and constructed two types microcosms with different initial counts of virus-like particles (VLPs) and sulfate content, namely SM, and SV microcosms. As shown in supplementary Fig. 12, approximately 500 mL of production water was filtered through sterile gauze to remove crude oil and obtain the fraction containing microbes and viruses. The filtrate was further divided into two parts, part of the filtrate was centrifuged (3500g at 4°C for 40 min) to obtain microbial pellets. Other part of filtrate was filtered through a 0.22-μm-pore-size polycarbonate membrane (45 mm diameter; Millipore, Bedford, MA, United States) to obtain the fractions water containing viruses. Afterward, part of 0.22 μm filtrate was filtered through tangential flow filtration equipment (TFF) with a 100 kDa tangential flow membrane package (Sartorius, VIVAFLOW 50 100,000 MWCO, Germany) to obtain virus-free production water. So far, basic components needed for microcosms were both acquired. The two type microcosms were set up using the above-mentioned basic components as described below. In brief, all microcosms were set up with sterilized serum bottles (internal volume 500 mL), containing 2 g sterile crude oil, 30 mL Brackish medium, and 300 mL water containing viruses or virus-free. Specifically, in SV microcosms, microbial pellets were diluted three times with virus-containing filtrate (Microbes+Viruses). Similarly, in SM microcosms, microbial pellets were diluted three times with virus-free filtrate (Microbes−Viruses). In addition, SV and SM microcosms were supplemented with sulfate to the final concentration were 4 g/L at the beginning of the culture. All treatments were set up in triplicates. After removing oxygen, all microcosms were incubated at 30°C in the dark. During the cultivation process, the number of sulfides was measured using total Sulfide Quantification Kit (LIANHUA, China). The detailed description of DNA extraction, library construction, sequencing, and statistical analysis of microcosms performed in this work can be found in the Supplemental Materials and methods section.

## Data availability

### Funding

#### Ethics approval

Not applicable.

#### Consent to participate

Not applicable.

#### Consent for publication

Not applicable.

