## supplemental Files for "Diversity and ecological functions of viruses inhabiting the oil reservoirs"

Supplementary material

**Supplementary Results and Discussion**

**AMGs encoded by viromes in oil reservoirs**

As shown in Supplementary Fig. 4c, d, for electron transport chain pathway, two AMGs involving the electron transport chain pathway were both found in one virus genomes, namely NADH-quinone oxidoreductase (respiratory complex I, NQOS, nuoA, nuoD, nuoH) and F-type H^+^-transporting ATPase subunit a (ATPF0A). NQOS is a central player in cellular energy metabolism and serves as a primary entry point for electrons from NADH into many respiratory chains^1-3^. F-type H^+^-transporting ATP synthases catalyze formation of ATP from ADP and inorganic phosphate at the expense of energy stored in the transmembrane gradient of protons^4^. ATP is the universal major energy source in a variety of relevant biological processes^5^. These results indicated that viruses may potentially intervene in the energy metabolism of host cells. For sulfur metabolism, AMGs encoding CysH were discovered in virus genome, which encodes phosphoadenosine phosphosulfate reductase, a key enzyme of the assimilatory sulfate reduction pathway^6^. For C1 metabolism, we discovered AMGs encoding pyruvate phosphate dikinase (PPDK) in virus genome, PPDK participates in the reductive tricarboxylic acid (rTCA) cycle^7^, and catalyzes the synthesis phosphoenolpyruvate from pyruvate^8^. In addition, AMGs encoding 2-oxoglutarate: ferredoxin oxidoreductase (KorA, KorB) were detected in samples, KorAB is a critical carbon-fixing enzyme in the rTCA cycle. KorAB catalyzes the reductive carboxylation of succinyl-CoA to 2-oxoglutarate of rTCA cycle^9^, another carboxylation reaction of rTCA cycle is the reductive carboxylation of 2-oxoglutarate to isocitrate, catalyzed by isocitrate dehydrogenase (IDH)^10,11^. In oil reservoirs, we found one AMGs encoding IDH, indicating that viruses may have the potential to mediate carbon fixation in host cells.

**Sulfur metabolism mediated by viruses in oil reservoirs**

We identified 484 host MAGs (39.77% of host MAGs) with potential sulfur metabolism ability and distributed in 73.38% samples based on the presence of sulfur metabolism functional marker genes within the host MAGs genome, these host MAGs were infected by 535 viruses, including 442 (82.62%) temperate viruses and 72 (13.46%) virulent viruses (Fig. 6a). Among these host MAGs, 50.00% host MAGs were predicted to participate in assimilatory sulfate reduction (ASR) in this process, sulfate is reduced to hydrogen sulfide and then included in cysteine and methionine biosynthesis, which is a structural block for proteins and polypeptides^12^. 18.80% of the host MAGs were predicted to participate in thiosulfate oxidation metabolism (TSO), which employs sulfur oxidation enzyme complex to oxidize thiosulfate and produce sulfate^13^. 6.82% of the host MAGs were predicted to participate in dissimilatory sulfate reduction (DSR), in this process, sulfate-reducing microorganisms reduce sulfate and produce toxic hydrogen sulfide for energy to grow^14,15^. 0.62% of the host MAGs were predicted to participate in sulfate oxidation (SO). In addition, 17.98% of the host MAGs were predicted to possess the ability for assimilatory sulfate reduction and thiosulfate oxidation, 0.83% of the host MAGs were predicted to possess the ability for assimilatory sulfate reduction and sulfate oxidation, 4.96% of the host MAGs were predicted to possess the ability for both thiosulfate oxidation and sulfate oxidation (Supplementary Fig. 7c).

**Evidence of viruses mediated sulfide production in microcosms**

In this study, we obtained a total of 2,037 putative viral contigs (with sizes > 10 kb). After clustering, a total of 786 vOTUs were retained. The genome quality of vOTUs consisted of 1.40% of high-quality, 13.87% of medium-quality, and 47.58% low quality vOTUs, while the quality of remaining 37.15% of vOTUs could not be determined. Based on the latest ICTV classification, the taxonomic analysis conducted with PhaGCN2.0 showed that 65.27% of vOTUs could not be annotated at the family level. The majority vOTUs were classified into the following families: Peduoviridae (20.36%), Casadabanvirus (2.80%), Triavirus (1.40%), and Herelleviridae (1.02%).

To study virus-host dynamics in greater depth, we reconstructed MAGs by binning shotgun metagenomic contigs. Overall, we successfully assembled 762 medium- to high-quality MAGs across all samples. In order to determine the association between MAGs and viruses, two different in silico methods were employed: CRISPR spacer linkages, and tRNA matching. Through these approaches, we found that 17.23% of viruses were linked to 218 predicted host MAGs. These predicted host MAGs spanned 3 archaeal and 17 bacterial phyla. Among the bacterial phyla, the top five predicted host MAGs were Desulfobacterota (63), Bacteroidota (35), Thermotogota (19), Actinobacteriota (16), and Patescibacteria (14) (Supplementary Fig. 8a,b). Additionally, we observed the presence of generalist viruses in both SM and SV microcosms, indicating a polyvalence nature. Furthermore, we found that viral and bacterial abundances correlated positively with each other (Supplementary Fig. 8e). Together, these results suggested that bacterial and viral community dynamics were tightly coupled within the microcosms.

**Supplemental methods**

### Detection of putative viral auxiliary metabolic genes (AMGs)

In order to identify putative viral auxiliary metabolic genes (AMGs) that may have a role in host metabolism during the infection cycle, we used DRAM-v v1.3.5^16^ to recover putative AMGs from vOTUs. Because DRAM-v requires VirSorter output, we re-ran all vOTUs through VirSorter v2.1. For DRAM-v output, as suggested previously^17-19^, to be conservative, we manually scanned the annotation output to improve the confidence in AMG identification, in particular, only putative AMGs with an auxiliary score < 4 were retained, and no viral flag (F), transposon flag (T), viral-like peptidase (P), or attachment flag (A) could be present, and putative AMGs that did not have a gene ID or a gene description were also discarded^17^. In addition, putative AMGs predicted to be involved in organic nitrogen, nucleotide metabolism, and predicted to be glycosyl transferases and ribosomal proteins were removed from downstream consideration, because some viruses can encode their own glycosyl transferases^20^. Moreover, to avoid false-positive results for selected AMGs caused by possible pollution of host sequences, we were looking for the presence of viral hallmark genes or virus-like genes in upstream and downstream of the putative AMG, only the putative AMGs located between or alongside two viral hallmark genes or virus-like genes were considered high-confidence viral AMGs for further analysis. Finally, genome maps for five vOTUs encoding AMGs were visualized based on DRAM-v and VirSorter2 annotations, protein structural homology searches were performed using the Phyre2 web portal^21^ using a confidence threshold of = 98%, coverage threshold of > 80%, and identity threshold of > 30%.

### Analysis of viral community

All statistical analyses were performed in R version 4.0.0^22^. The world map was created with the maps library and created using the geom_polygon function in “ggplot2” package. All histogram plots were created using the geom_histogram function in “ggplot2” package, all bar plots were created using the geom_bar function in “ggplot2” package, all pie plots were created using the pie function. Alpha and beta diversity of viral communities were calculated using “vegan” v2.5-7 package^22,23^. Nonmetric multidimensional scaling (NMDS) was conducted based on Bray-Curtis dissimilarities generated from vOTUs tables with viral abundances (RPKM) using the “vegdist” function. To further determine the significant difference of viral community composition between different continents, a similarity analysis (ANOSIM) was performed using the “anosim” function. In addition, to understand distance-decay relationships (DDRs) of viral community, pairwise geographic distances between samples were calculated from the latitude and longitude coordinates using the “geosphere” library, and the relationships between geographic distances and viral community similarities (1 − dissimilarity of the Bray-Curtis metric) were calculated by ordinary least-squares regressions.

To determine the contribution of different ecological processes to community assembly, null model analysis was carried out using the framework described by Stegen et al^24^. The null model expectation was generated using 999 randomizations in R. Two metrics, including β-nearest taxon index (βNTI) and Bray-Curtis-based Raup-Crick (RC_Bray_), were calculated to divide community assembly into five processes, namely, homogeneous selection, variable selection, dispersal limitation, homogeneous dispersal, and drift. βNTI > 2 indicates heterogeneous selection, βNTI < −2 indicates homogeneous selection. |βNTI| < 2 and RC_Bray_ < −0.95 indicate homogenizing dispersal, |βNTI| < 2 and RC_Bray_ > 0.95 indicate dispersal limitation, |βNTI| < 2 and |RC_Bray_| < 0.95 indicate drift assembly^24^. Heatmaps were created using the geom_raster function in “ggplot2” package.

Pearson correlations were performed using ‘quickcor’ function in “ggcor” package to assess the relationships between the biotic and abiotic variables. Mantel tests were performed to using ‘mantel_test’ function in “vegan” package to reveal the correlations between community structures (Bray-Curtis distance) and biotic and abiotic variables.

### DNA extraction, library construction, and sequencing of Microcosms

After 160 days (T1) and 570 days (T2) of incubation, 330 mL cultures were filtered by 0.22-μm-pore-size polycarbonate membranes (45 mm diameter; Millipore, Bedford, MA, United States) to obtain microbial cells. Total microbial DNA was extracted and sequencing as described before. Metagenomes of all microcosms were analyzed as described before.

### Statistical analyses of microcosm experiments

All statistical analyses were performed in R version 4.0.0^22,23^. The difference analysis of H_2_S between different microcosms was calculated using two-tailed T-test. The alpha diversity metrics, including richness, and Shannon (Shannon-Wiener diversity), were calculated using the ‘vegan’ and ‘picante’ packages. For beta diversity, the principal components analysis (PCoA) based on Bray-Curtis distance and a similarity analysis (ANOSIM) were performed in the ‘vegan’ package. The differential enrichment KEGG modules related to energy metabolism and hydrocarbon degradation were identified according to their reporter scores and reporter features. The final set of differential enrichment KEGG modules was determined as the union of the KEGG modules based on reporter scores and reporter features. Reporter scores and reporter features were calculated as described before ^25,26^. To understand the significant change of gene abundance between different microcosms, KEGG homologs were combined and summed according to gene clusters table of non-redundant gene sets. Then we conducted difference analysis for KO using two-tailed T-test.

**Supplemental figures**

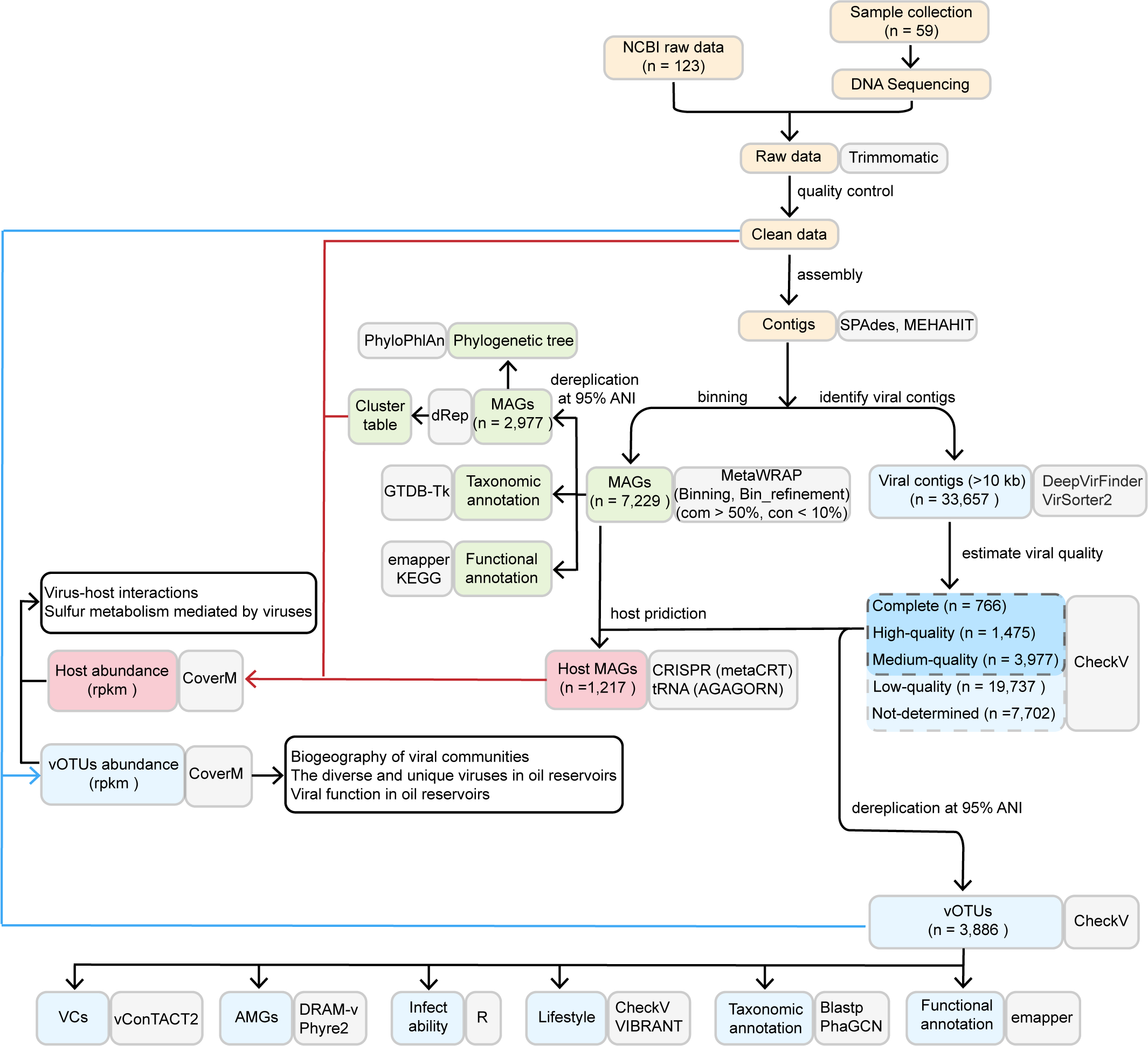

Supplementary Fig. 1 The diagram showing the bioinformatic workflow for the identification of viruses. Yellow: data collection and initial processing; green: microbial analysis; blue: viral analysis; red: interactions between hosts and viruses; grey: software used.

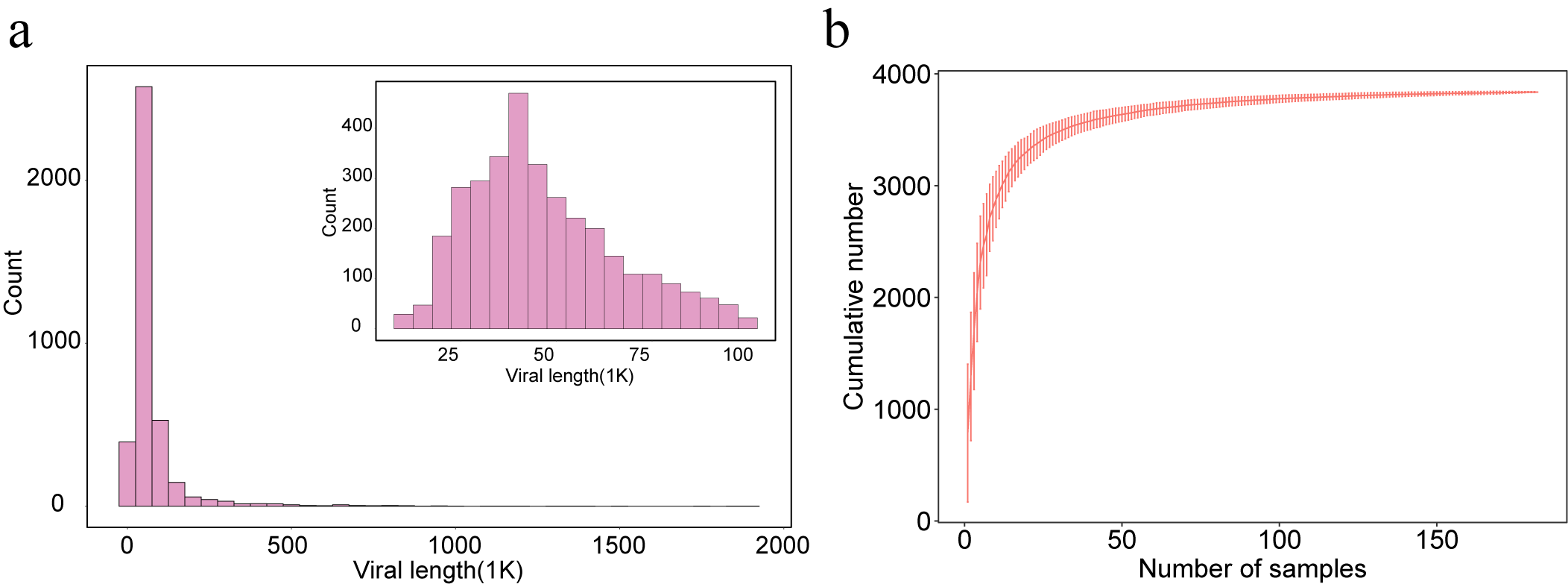

Supplementary Fig. 2

a, Histogram illustrating the genome size distribution of vOTUs (≥ 10 kb).

b, Accumulation curve of detected vOTUs numbers from all 182 samples. Dots represent the average number of vOTUs for all combinations of a given number of samples, and error bars represent the range.

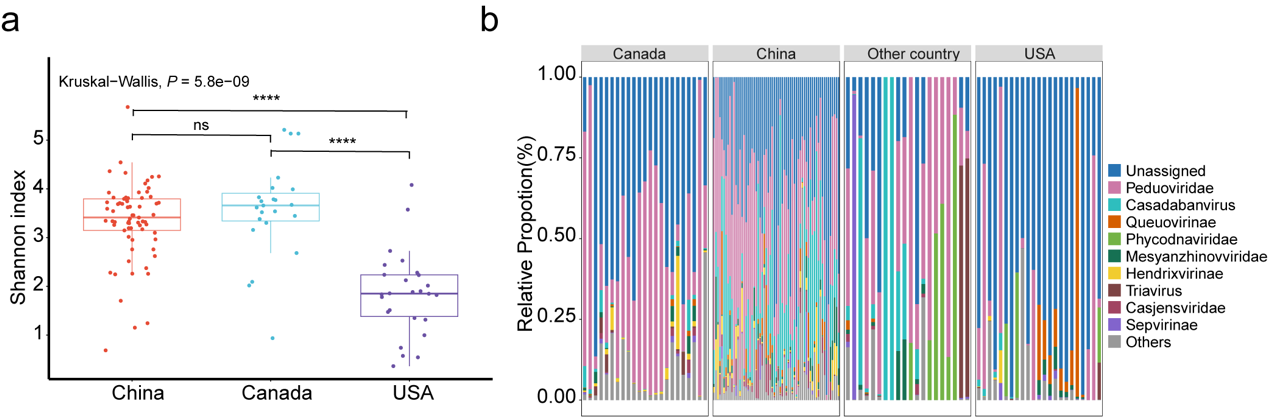

Supplementary Fig. 3 Viral diversity (a) and composition (b) of oil reservoir samples collected from different countries. Asterisks denote significance, with **** indicating *P* < 0.001.

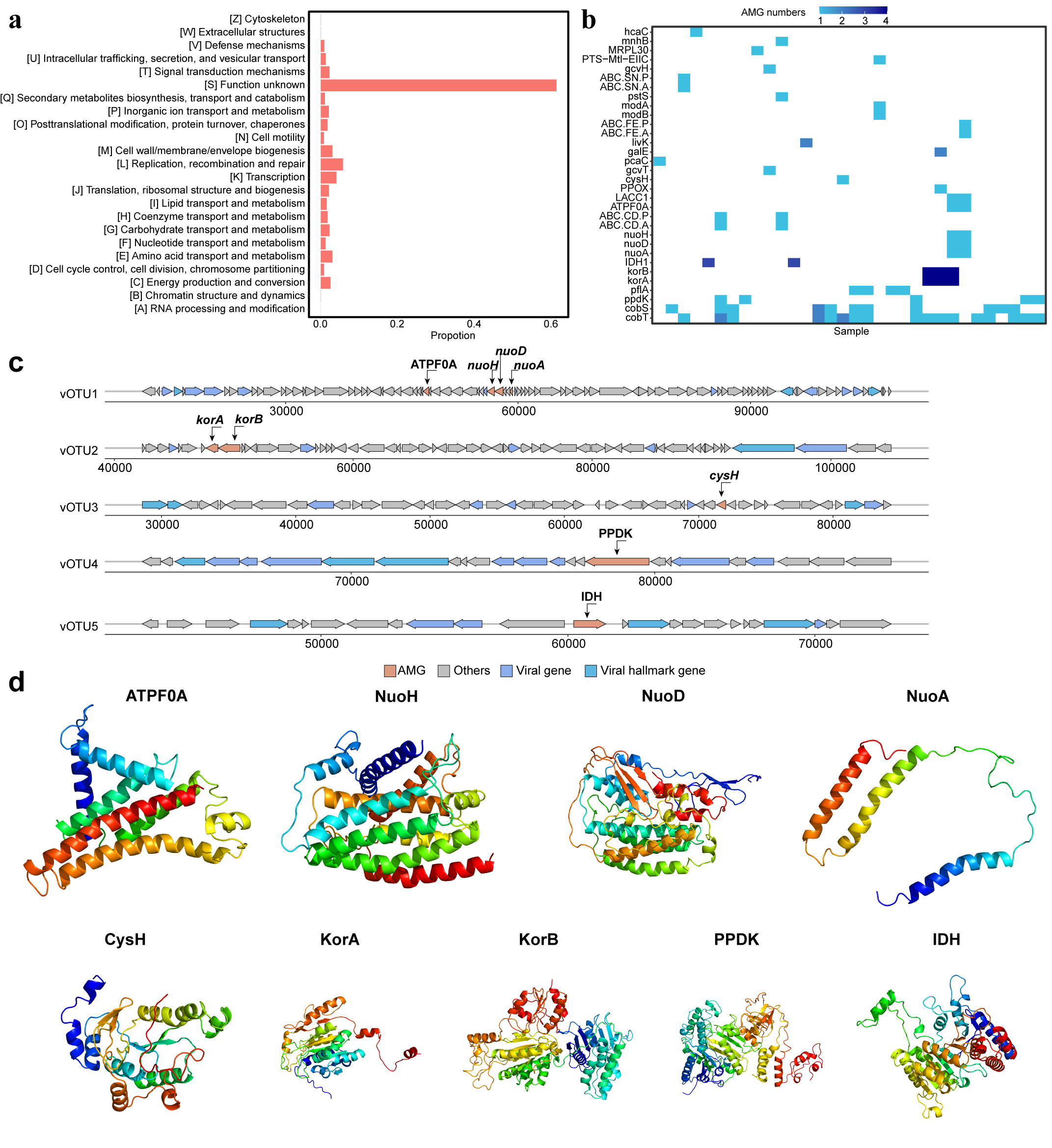

Supplementary Fig. 4 Functional genes encoded by viromes in oil reservoirs. a, Relative abundances of vOTUs functions in the oil reservoirs as annotated by eggNOG v5.0.0 database. b, The distribution of different metabolism categories of virus-encoded auxiliary metabolic genes (AMGs) of vOTUs in oil reservoirs samples. c, Genome map of representative AMG-encoding viruses predicted to participate in energy metabolism, AMGs are in orange, virus-like genes are in blue, viral hallmark genes are in dark blue, and non-virus-like or uncharacterized genes are in gray. d. Tertiary structures of AMGs predicted to participate in energy metabolism based on structural modelling using Phyre2.

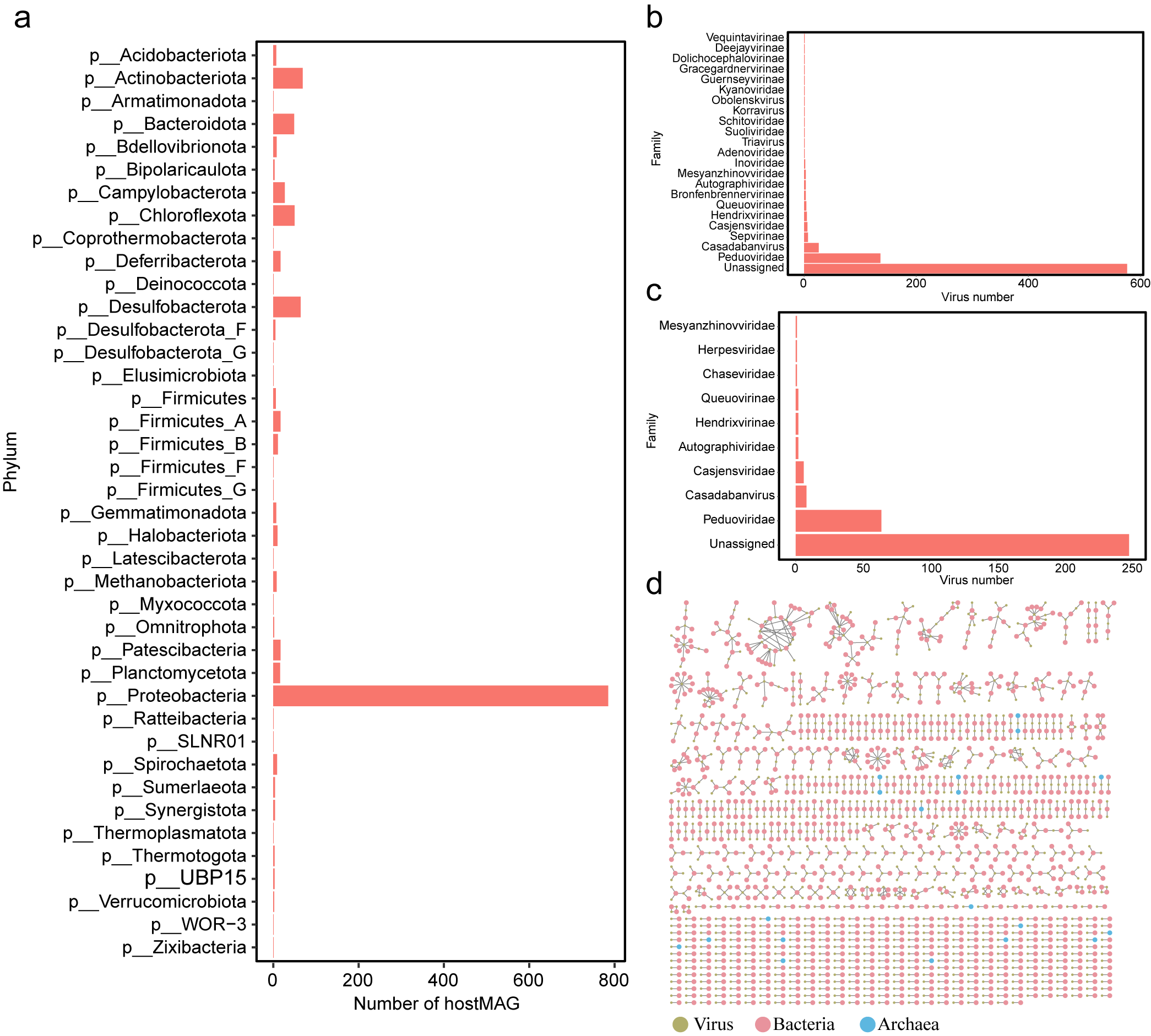

Supplementary Fig. 5

a, Histogram showing the taxonomic distribution of predicted host MAGs at phylum level. b, c, Histogram showing the taxonomic distribution of specialist (b) and generalist (c) viruses. d. Interactions between host MAGs and viruses. Nodes represent viruses (green) and host MAGs (bacteria: blue; archaea: pink), and edges represent the relationship between viruses and host MAGs based on CRISPR spacers and tRNA sequences.

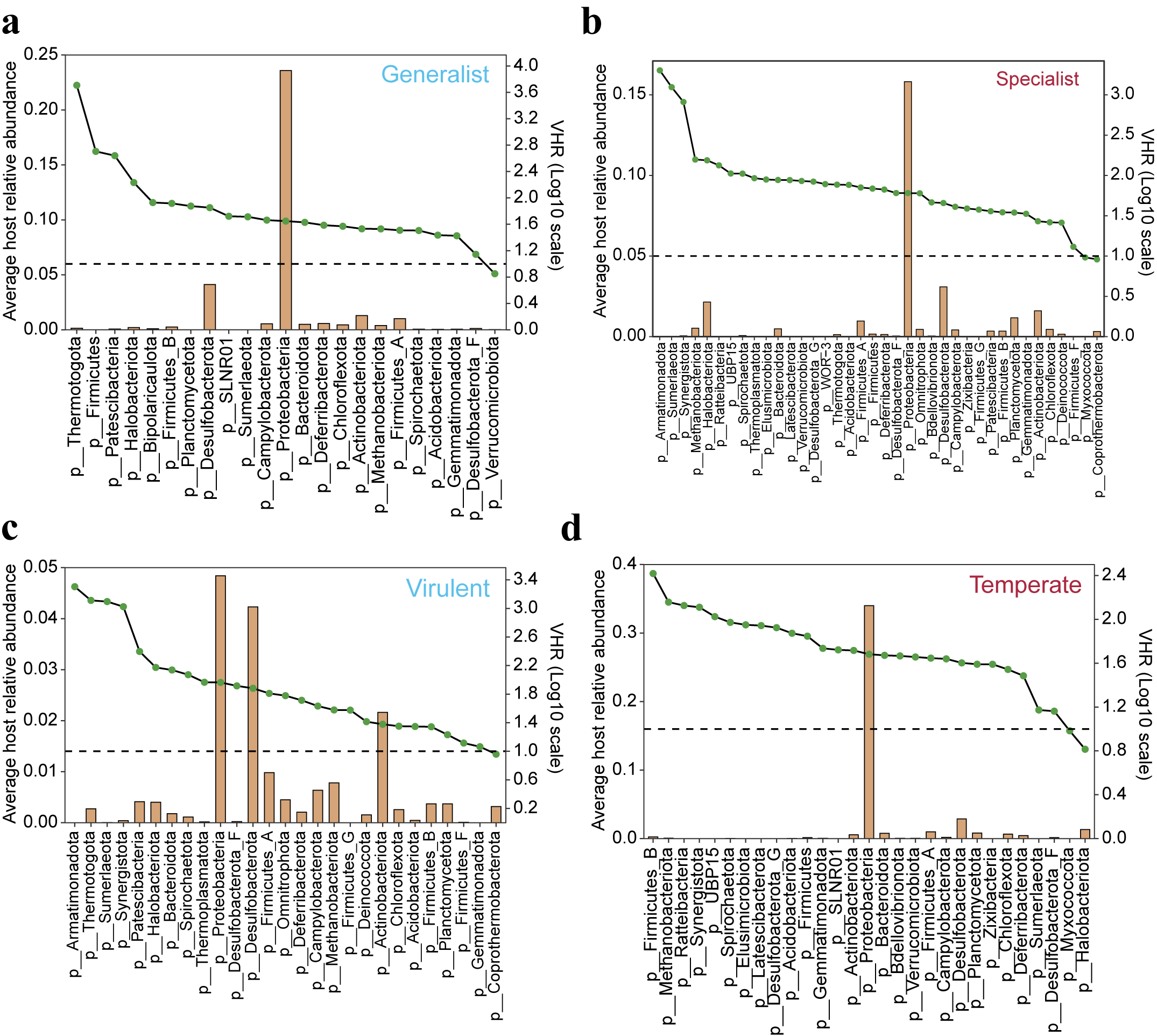

Supplementary Fig. 6 Lineage-specific generalist (a) or specialist (b) or virulent (c) or temperate (d) virus-host abundance ratios (VHRs) for all predicted microbial hosts. The bar graph represents the relative abundance of microbial hosts at the phylum level, the green dot represents the VHR, and the red vertical line represents the 1:1 ratio.

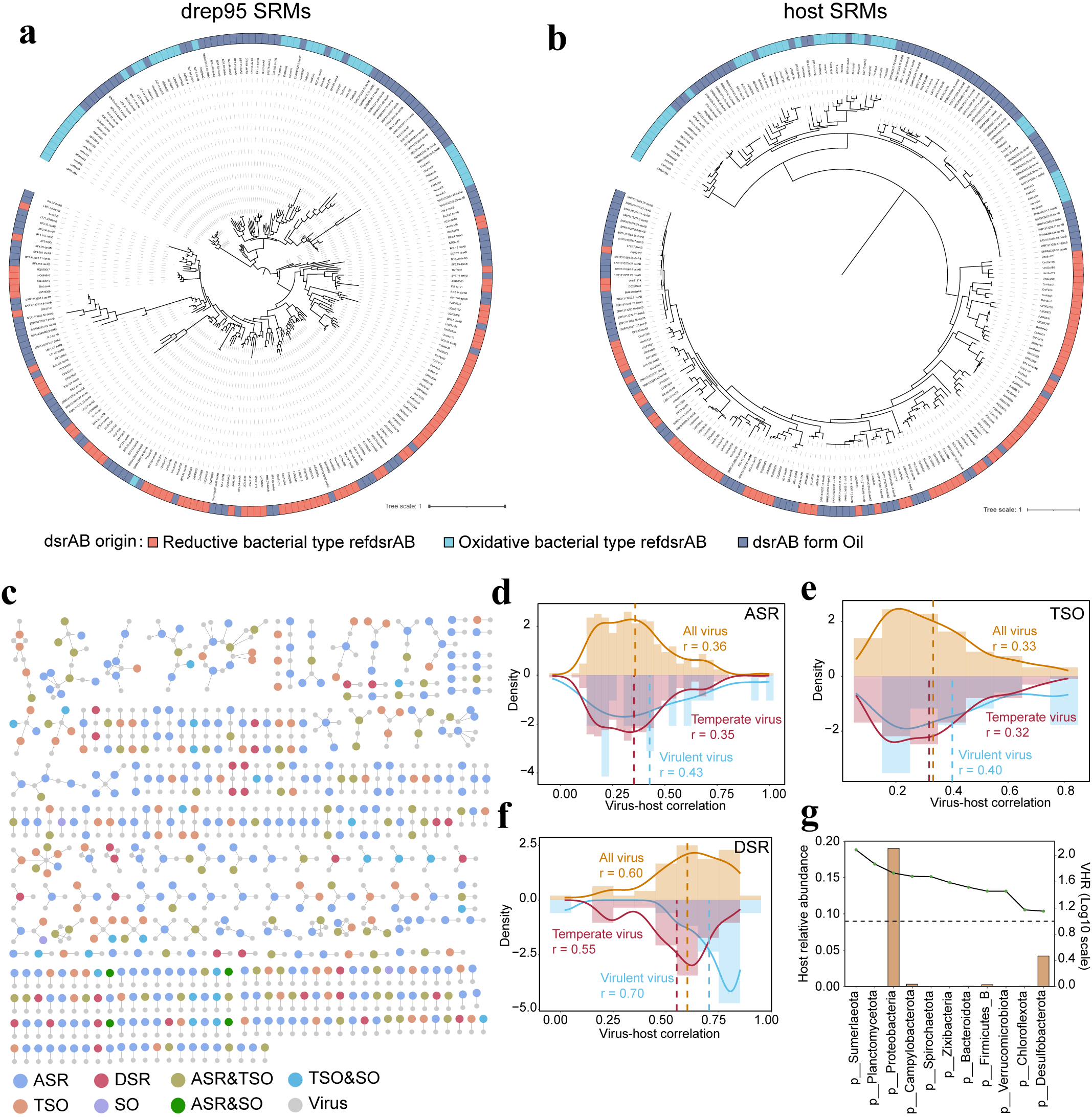

Supplementary Fig. 7 Close interactions between the oil reservoir viruses and host MAGs with potential sulfur metabolism ability. a, b, Phylogenic analysis of dissimilatory sulfite reductases DsrAB from all non-redundant MAGs (a) and host MAGs (b). Concatenated DsrAB protein sequences identified in this study are marked in grey color. Assignment of oxidative or reductive type DsrAB was according to previous studies^27,28^. Bootstrap values were based on 1000 replicates. c, Interactions between host MAGs with potential sulfur metabolism ability and viruses. Grey nodes represent viruses and other color nodes represent host MAGs with different potential sulfur metabolism ability, and edges represent the relationship between viruses and host MAGs with potential sulfur metabolism ability based on CRISPR spacers and tRNA sequences. d, e, f, Distribution of viruses and hosts with different sulfur metabolism ability correlations. Orange, blue, and red colors represent the distributions of all viruses, virulent viruses, and temperate viruses, respectively. Dashed lines show the average correlation in the distribution. g, Lineage-specific virus-host abundance ratios (VHRs) for all predicted microbial hosts. The bar graph represents the relative abundance of microbial hosts at the phylum level, the green dot represents the VHR, and the black vertical line represents the 1:1 ratio.
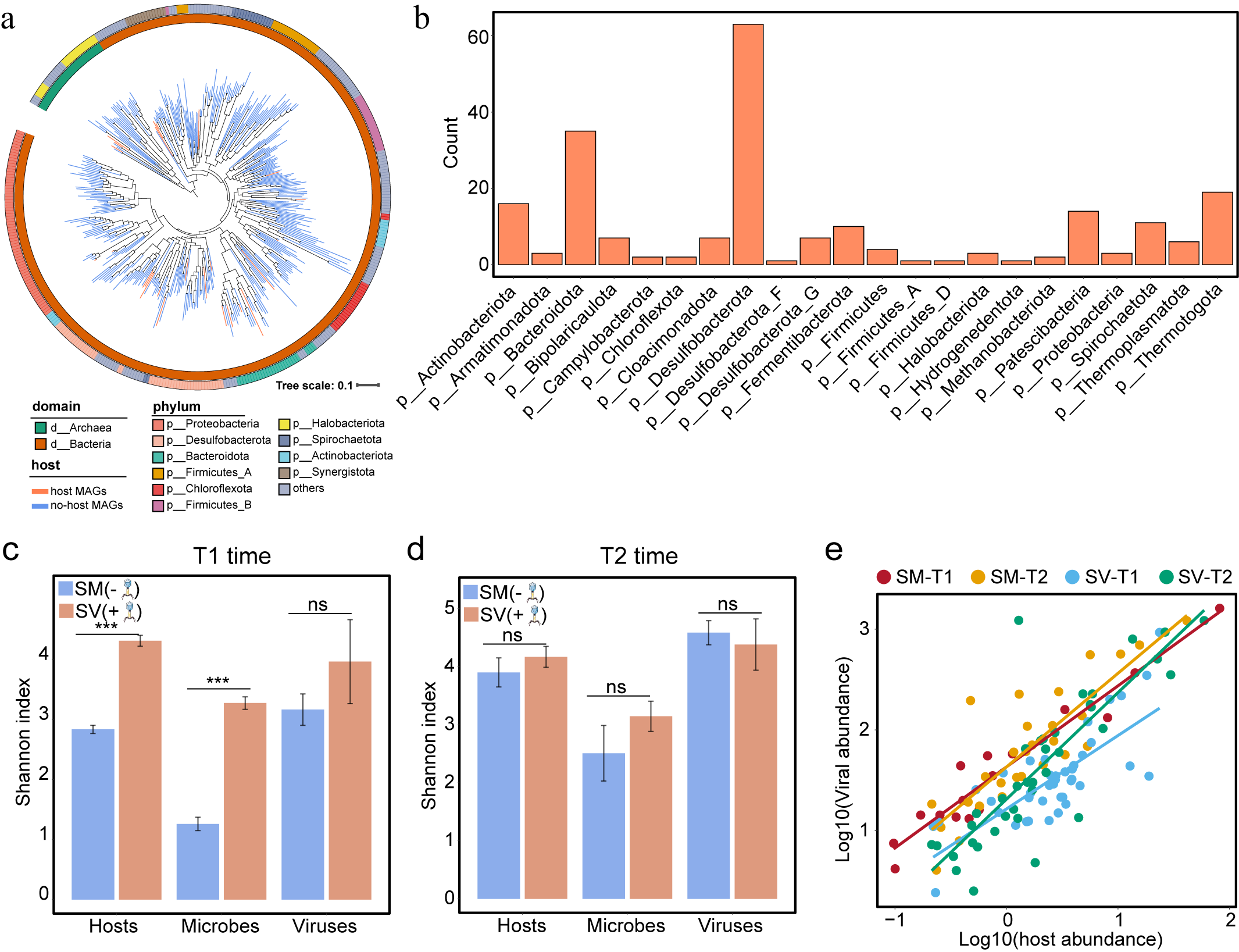

Supplementary Fig. 8 Virus-host linkages and the dynamics of host MAGs and viruses in microcosms. a, Maximum-likelihood phylogenetic trees of MAGs at the phylum level detected in microcosms. The color of clades indicates whether MAGs are host MAGs (Orange represents host MAGs, blue represents non-host MAGs). The inner circle represents domain and the outer circle represents phylum annotated by GTDB. The tree was constructed by PhyloPhlAn and visualized by iTOL. b, Histogram showing the taxonomic distribution of predicted host MAGs at phylum level. c, d. Comparison of the Shannon index of the host sub-community, microbial community, and viral community among SM and SV microcosms at T1 (c) and T2 (d) sampling time points. e, Significant positive correlation between the relative abundance of viruses and their hosts in the microcosms.

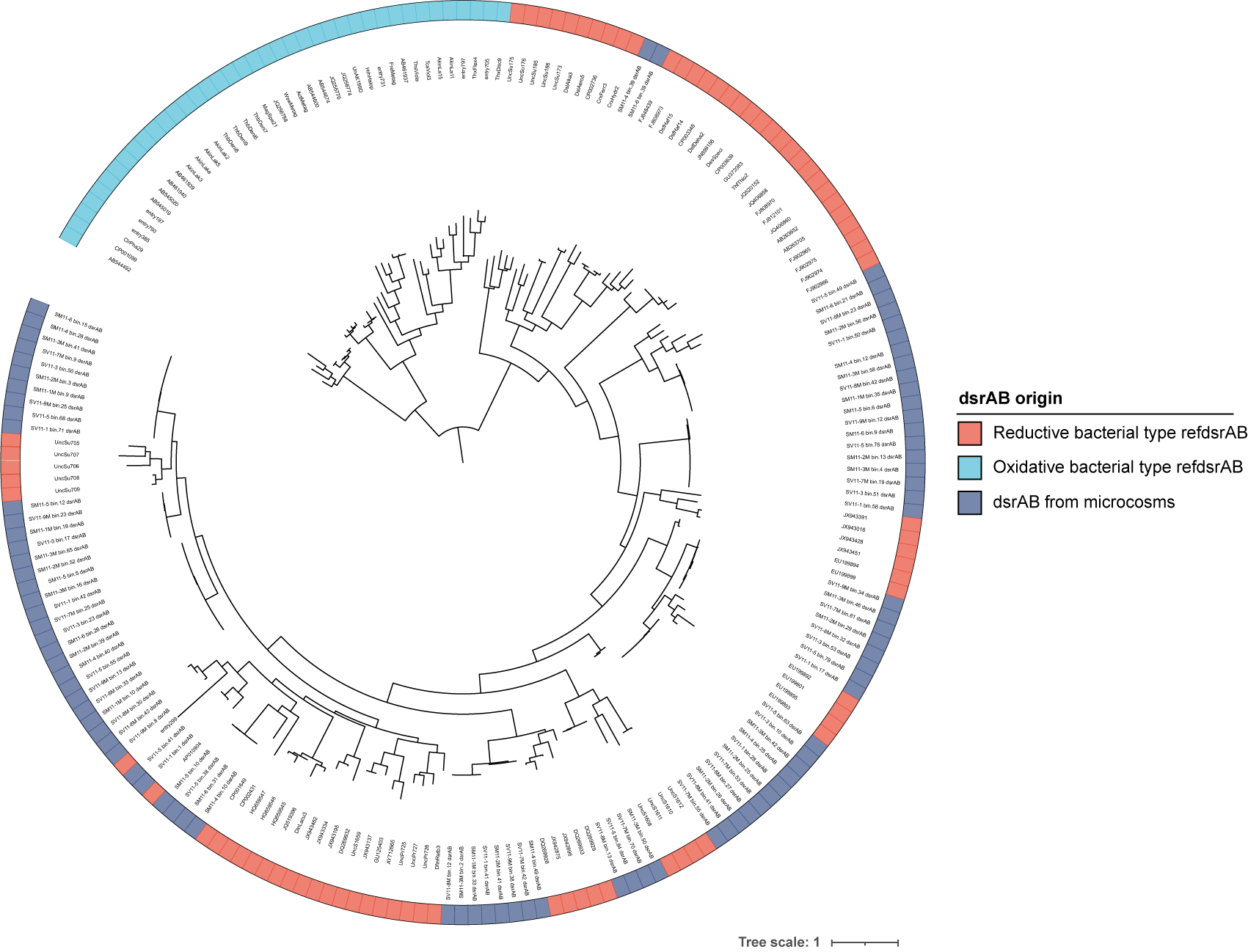

Supplementary Fig. 9 Phylogenic analysis of dissimilatory sulfite reductases DsrAB in all MAGs from microcosms. Concatenated DsrAB protein sequences identified in this study are marked in grey color. Assignment of oxidative or reductive type DsrAB was according to previous studies^27,28^. Bootstrap values were based on 1000 replicates.

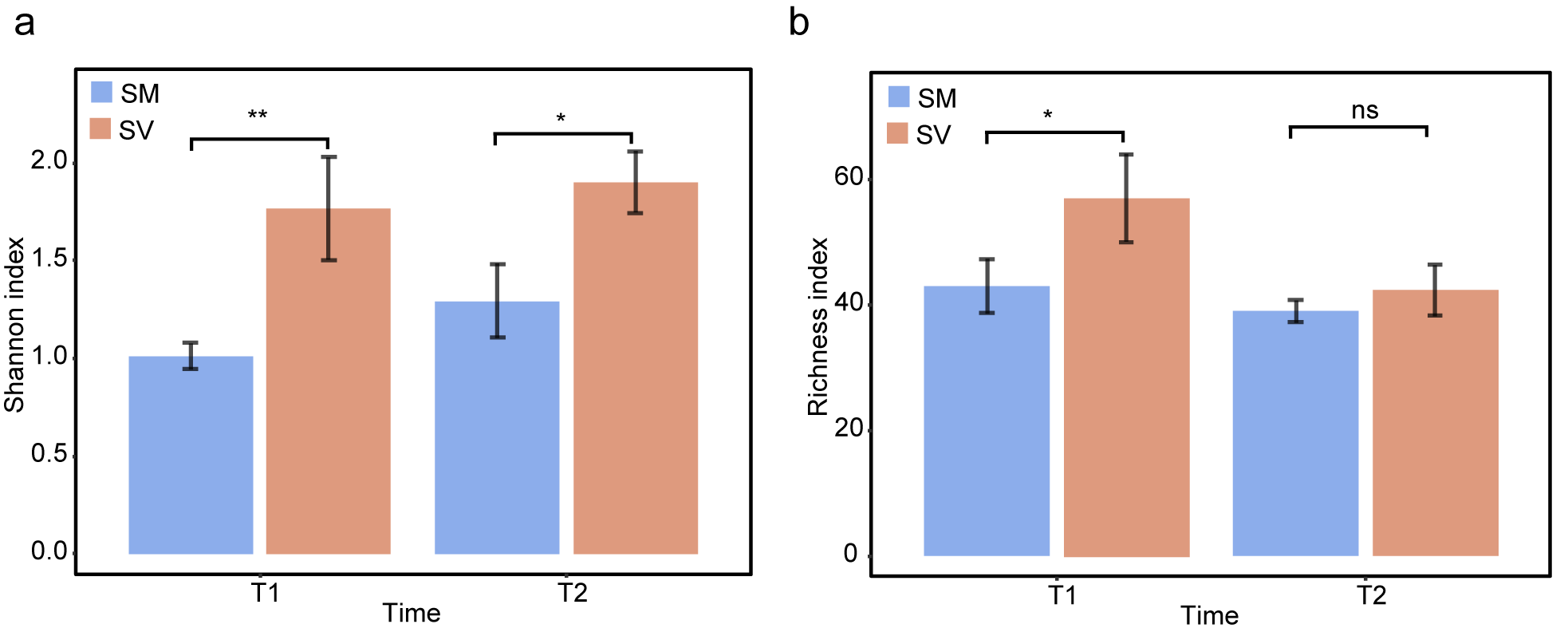

Supplementary Fig. 10 Comparison of the Shannon (a) and richness (b) index of the microbial community among SM and SV microcosms at T1 and T2 sampling time points.

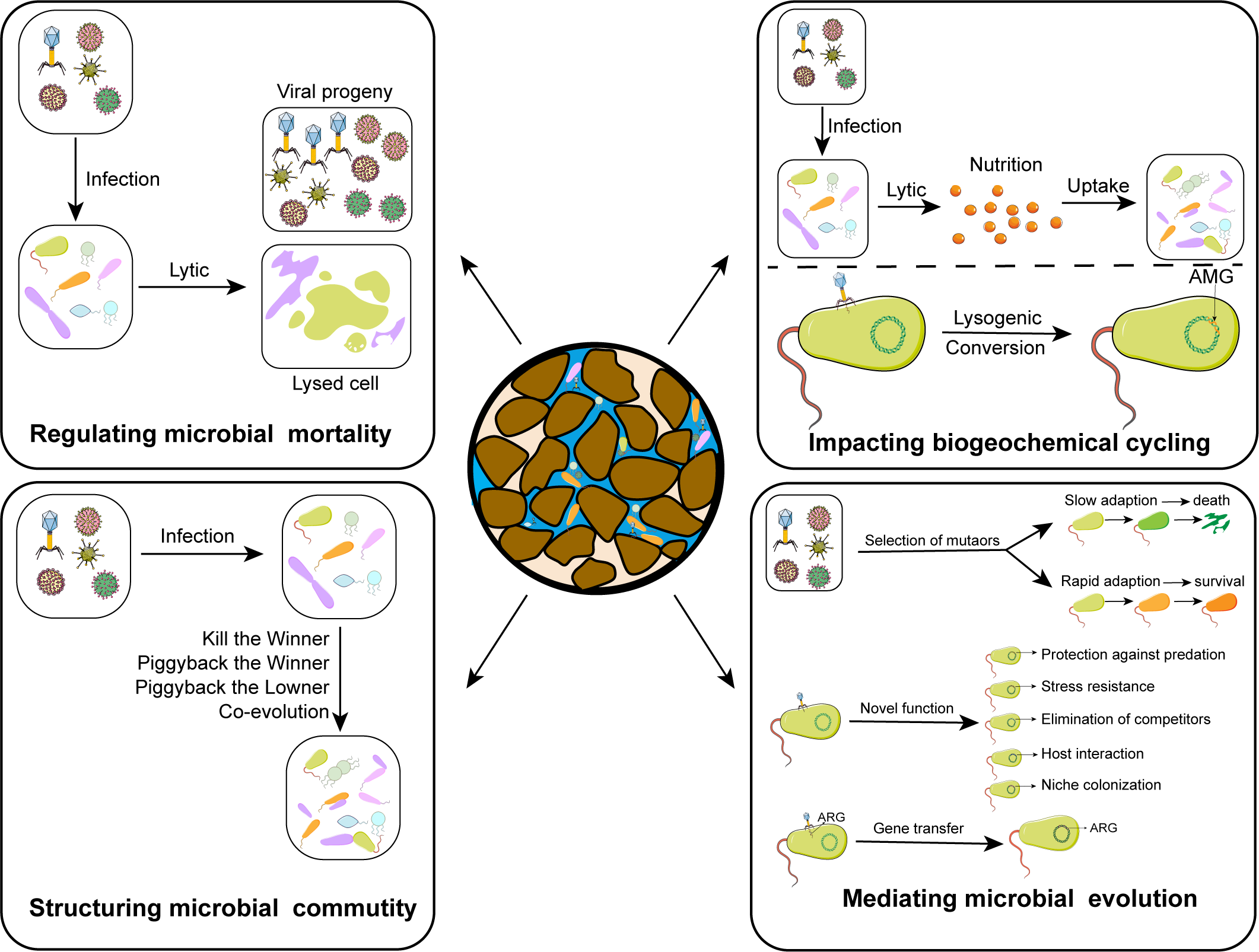

Supplementary Fig. 11 Overview of potential ecological roles of viruses in oil reservoirs.

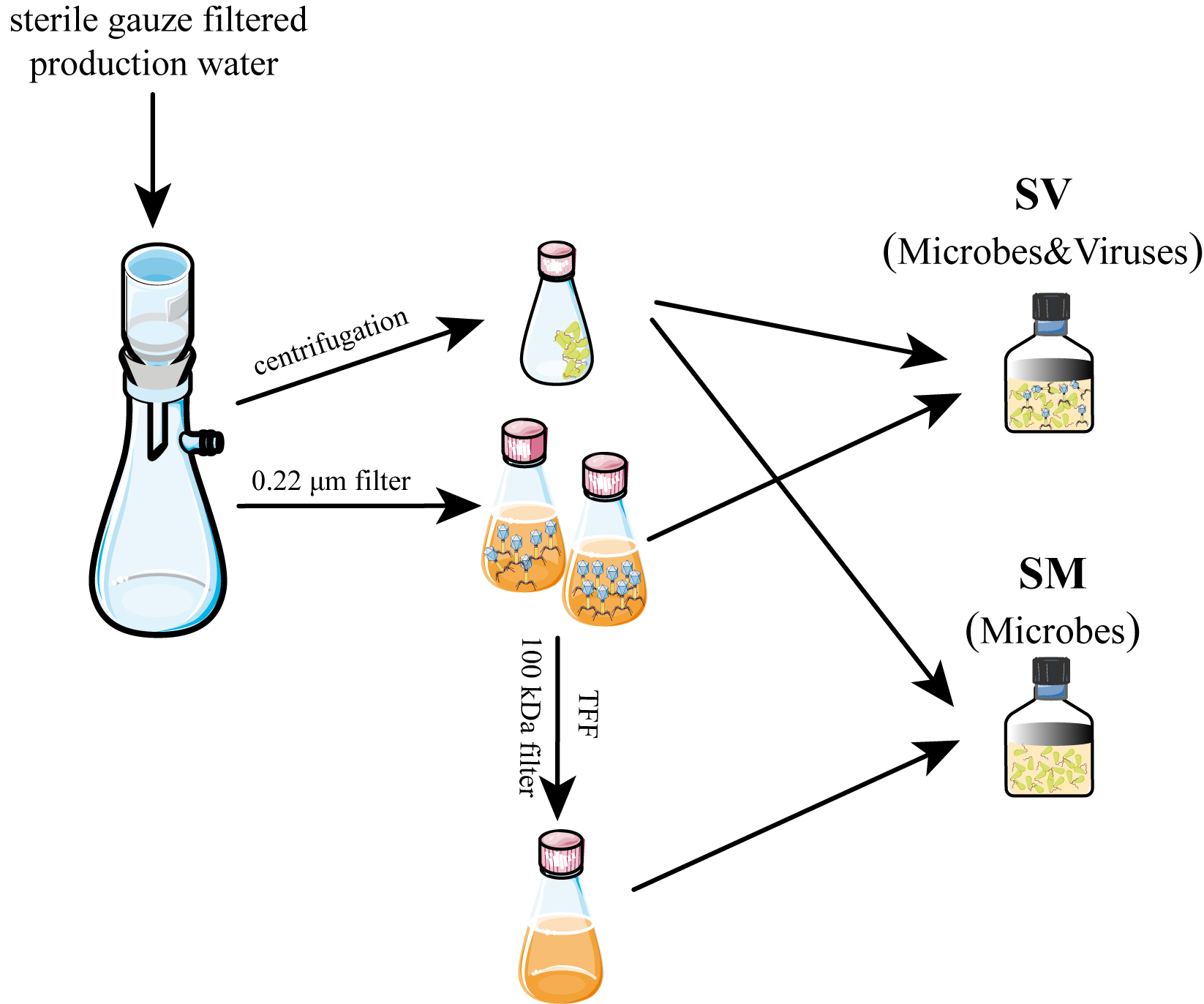

Supplementary Fig. 12 Schematic of experimental setup used in this study.

1	Kaila, V. R. I. Resolving Chemical Dynamics in Biological Energy Conversion: Long-Range Proton-Coupled Electron Transfer in Respiratory Complex I. *Accounts of chemical research* **54**, 4462-4473, doi:10.1021/acs.accounts.1c00524 (2021).

2	Parey, K., Wirth, C., Vonck, J. & Zickermann, V. Respiratory complex I - structure, mechanism and evolution. *Current opinion in structural biology* **63**, 1-9, doi:10.1016/j.sbi.2020.01.004 (2020).

3	Hoeser, F., Weiß, M. & Friedrich, T. The clinically relevant triple mutation in the mtND1 gene inactivates Escherichia coli complex I. **596**, 1124-1132, doi:https://doi.org/10.1002/1873-3468.14325 (2022).

4	Pierson, H. E., Uhlemann, E.-M. E. & Dmitriev, O. Y. Interaction with Monomeric Subunit c Drives Insertion of ATP Synthase Subunit a into the Membrane and Primes a-c Complex Formation*. *Journal of Biological Chemistry* **286**, 38583-38591, doi:https://doi.org/10.1074/jbc.M111.294868 (2011).

5	Vu Huu, K., Zangl, R., Hoffmann, J., Just, A. & Morgner, N. Bacterial F-type ATP synthases follow a well-choreographed assembly pathway. *Nat Commun* **13**, 1218, doi:10.1038/s41467-022-28828-1 (2022).

6	Mara, P. *et al.* Viral elements and their potential influence on microbial processes along the permanently stratified Cariaco Basin redoxcline. *The ISME Journal* **14**, 3079-3092, doi:10.1038/s41396-020-00739-3 (2020).

7	Hügler, M. & Sievert, S. M. Beyond the Calvin Cycle: Autotrophic Carbon Fixation in the Ocean. **3**, 261-289, doi:10.1146/annurev-marine-120709-142712 (2011).

8	Imaizumi, N. *et al.* Characterization of the gene for pyruvate,orthophosphate dikinase from rice, a C3 plant, and a comparison of structure and expression between C3 and C4 genes for this protein. *Plant molecular biology* **34**, 701-716, doi:10.1023/a:1005884515840 (1997).

9	Kim, J., Oh, E. K., Kim, E. J. & Lee, J. K. Photoautotrophic Growth Rate Enhancement of Synechocystis sp. PCC6803 by Heterologous Production of 2-Oxoglutarate:Ferredoxin Oxidoreductase from Chlorobaculum tepidum. *Biology* **12**, doi:10.3390/biology12010059 (2022).

10	Hügler, M. & Sievert, S. M. Beyond the Calvin cycle: autotrophic carbon fixation in the ocean. *Annual review of marine science* **3**, 261-289, doi:10.1146/annurev-marine-120709-142712 (2011).

11	Kanao, T., Kawamura, M., Fukui, T., Atomi, H. & Imanaka, T. Characterization of isocitrate dehydrogenase from the green sulfur bacterium Chlorobium limicola. A carbon dioxide-fixing enzyme in the reductive tricarboxylic acid cycle. *Eur J Biochem* **269**, 1926-1931, doi:10.1046/j.1432-1033.2002.02849.x (2002).

12	Schiff, J. A. Pathways of assimilatory sulphate reduction in plants and microorganisms. *Ciba Foundation symposium*, 49-69, doi:10.1002/9780470720554.ch4 (1979).

13	Koch, T. & Dahl, C. A novel bacterial sulfur oxidation pathway provides a new link between the cycles of organic and inorganic sulfur compounds. *The ISME Journal* **12**, 2479-2491, doi:10.1038/s41396-018-0209-7 (2018).

14	Rabus, R. *et al.* A Post-Genomic View of the Ecophysiology, Catabolism and Biotechnological Relevance of Sulphate-Reducing Prokaryotes. *Advances in microbial physiology* **66**, 55-321, doi:10.1016/bs.ampbs.2015.05.002 (2015).

15	Qian, Z., Tianwei, H., Mackey, H. R., van Loosdrecht, M. C. M. & Guanghao, C. Recent advances in dissimilatory sulfate reduction: From metabolic study to application. *Water Research* **150**, 162-181, doi:https://doi.org/10.1016/j.watres.2018.11.018 (2019).

16	Shaffer, M. *et al.* DRAM for distilling microbial metabolism to automate the curation of microbiome function. *Nucleic Acids Res* **48**, 8883-8900, doi:10.1093/nar/gkaa621 (2020).

17	Ter Horst, A. M. *et al.* Minnesota peat viromes reveal terrestrial and aquatic niche partitioning for local and global viral populations. *Microbiome* **9**, 233, doi:10.1186/s40168-021-01156-0 (2021).

18	Li, Z. *et al.* Deep sea sediments associated with cold seeps are a subsurface reservoir of viral diversity. *The ISME Journal* **15**, 2366-2378, doi:10.1038/s41396-021-00932-y (2021).

19	Zhao, J. *et al.* Novel Viral Communities Potentially Assisting in Carbon, Nitrogen, and Sulfur Metabolism in the Upper Slope Sediments of Mariana Trench. *mSystems* **7**, e0135821, doi:10.1128/msystems.01358-21 (2022).

20	Xiang, Y. *et al.* Crystal structure of a virus-encoded putative glycosyltransferase. *Journal of virology* **84**, 12265-12273, doi:10.1128/jvi.01303-10 (2010).

21	Kelley, L. A., Mezulis, S., Yates, C. M., Wass, M. N. & Sternberg, M. J. The Phyre2 web portal for protein modeling, prediction and analysis. *Nature protocols* **10**, 845-858, doi:10.1038/nprot.2015.053 (2015).

22	Ihaka, R. & Gentleman, R. R: A Language for Data Analysis and Graphics. *Journal of Computational and Graphical Statistics* **5**, 299-314, doi:10.2307/1390807 (1996).

23	Oksanen J, B. F., Kindt R, Legendre P, Minchin PR, O’Hara R, . Vegan: community ecology package. R package version 22-1. (2015).

24	Stegen, J. C. *et al.* Quantifying community assembly processes and identifying features that impose them. *The ISME Journal* **7**, 2069-2079, doi:10.1038/ismej.2013.93 (2013).

25	Feng, Q. *et al.* Gut microbiome development along the colorectal adenoma-carcinoma sequence. *Nat Commun* **6**, 6528, doi:10.1038/ncomms7528 (2015).

26	Oliveira, A. P., Patil, K. R. & Nielsen, J. Architecture of transcriptional regulatory circuits is knitted over the topology of bio-molecular interaction networks. *BMC systems biology* **2**, 17, doi:10.1186/1752-0509-2-17 (2008).

27	Müller, A. L., Kjeldsen, K. U., Rattei, T., Pester, M. & Loy, A. Phylogenetic and environmental diversity of DsrAB-type dissimilatory (bi)sulfite reductases. *Isme j* **9**, 1152-1165, doi:10.1038/ismej.2014.208 (2015).

28	Anantharaman, K. *et al.* Expanded diversity of microbial groups that shape the dissimilatory sulfur cycle. *Isme j* **12**, 1715-1728, doi:10.1038/s41396-018-0078-0 (2018).
